# Genomic, spatial, and evolutionary insights into a dominant *Mycoplasmatota* symbiont colonizing the body wall of deep-sea holothurians

**DOI:** 10.64898/2026.08.10.742674

**Authors:** Yu Yoshida, Yosuke Nishimura, Hajime Itoh, Masumi Hasegawa-Takano, Tsuyoshi Takano, Naohisa Wada, Kento Tominaga, Akito Ogawa, Wataru Iwasaki, Yasuhiro Gotoh, Takehiko Itoh, Tetsuya Hayashi, Susumu Yoshizawa

**Affiliations:** Graduate School of Frontier Sciences, The University of Tokyo; Institute for Extra-cutting-edge Science and Technology Avant-garde Research of Life (X-star), Japan Agency for Marine-Earth Science and Technology (JAMSTEC); Atmosphere and Ocean Research Institute, The University of Tokyo; Advanced Institute for Marine Ecosystem Change (WPI-AIMEC), Japan Agency for Marine-Earth Science and Technology (JAMSTEC); Meguro Parasitological Museum; Okinawa Institute of Science and Technology Graduate University (OIST); Center for Molecular Biodiversity Research, National Museum of Nature and Science; Department of Bacteriology, Faculty of Medical Sciences, Kyushu University; School of Life Science and Technology, Institute of Science Tokyo

**Author notes:** Correspondence: Yosuke Nishimura and Susumu Yoshizawa. These authors contributed equally to this work (Yu Yoshida, Yosuke Nishimura).

## Abstract

Subcuticular bacteria (SCB) are widespread symbionts of echinoderms and often dominate the body-wall microbiome, suggesting important roles in host physiology. However, their diversity, metabolic properties, and host associations remain poorly characterized. Here, we report a novel dominant SCB lineage associated with deep-sea holothurians, *Scotoplanes* spp. collected from the Northwest Pacific. We recovered two high-quality genomes, including a 649-kb complete circular genome, and propose a new genus and species, “*Candidatus* Abyssoplasma scotoplanesicola”, within *Mycoplasmatota*. The two genomes showed a highly reduced metabolic repertoire, lacking central pathways including glycolysis. In contrast, acidic cell-surface-associated proteins, including large proteins exceeding 5,000 amino acids, accounted for 27.6% of the complete genome and clustered near defense islands. Localized genome plasticity in these regions, revealed by comparison between the two closely related genomes, suggests a possible mechanism for diversification of cell-surface proteins at the host-symbiont interface. “*Candidatus* Abyssoplasma scotoplanesicola” occupied 76.4–98.9% of the body-wall microbiome of the *Scotoplanes* specimens. Fluorescence in situ hybridization analysis confirmed that these bacteria formed aggregates on the epidermal side of the body wall. Overall, this study provides genome-and spatially resolved views of dominant SCB in holothurians and offers evolutionary insights into host-interface diversification in the deep-sea holothurian body wall.

---

Keywords: Subcuticular bacteria; *Scotoplanes*; *Mycoplasmatota*; deep-sea symbiosis; comparative genomics; long CDSs;

## Introduction

Symbiotic relationships between animals and bacteria are widespread in marine ecosystems and play fundamental roles in host nutrition, immunity, development, and ecological adaptation [1–3]. In many well-studied systems, specific bacterial lineages dominate certain host tissues and perform essential physiological functions, as exemplified by bioluminescent and chemosynthetic symbioses [4–7]. Such tissue-restricted and highly dominant symbionts are often tightly integrated into host biology.

Echinodermata is a marine-restricted animal phylum in which bacterial symbionts have been implicated in diverse physiological functions, ranging from amino acid uptake through the body surface to digestive processing and morphological modulation [8–10]. Echinoderms uniquely host symbionts known as subcuticular bacteria (SCB) under the cuticular layer across all five extant classes [11,12]. These bacteria have been reported from 50–60% of echinoderm species examined in previous studies [11,13]. Bacterial communities associated with echinoderm body walls appear to exhibit high host specificity, and this tendency is even stronger in SCB [14–16]. SCB have been observed at high cell densities in brittle stars [17]. Taken together, these findings suggest that SCB are common in echinoderms and represent an important bacterial group due to their close host associations and high abundance. However, despite their potential importance, a genome-resolved understanding of SCB remains limited, constraining our understanding of their metabolic capabilities, interactions with hosts, and colonization strategies. Filling this gap is particularly important for deposit-feeding holothurians (sea cucumbers), including deep-sea holothurians of the genus *Scotoplanes* (Elasipodida: Elpidiidae), commonly known as sea pigs. *Scotoplanes* spp. are abundant deposit feeders widely distributed in the deep sea [18,19] and have been shown to selectively ingest recently settled organic-rich particles, thereby contributing substantially to the processing of organic particle inputs on the seafloor [20].

To date, only two SCB genomes have been reported, each from a different echinoderm class. In the crown-of-thorns starfish, a dominant *Spirochaete* SCB is shared across geographically distant host populations, and its complete genome from metagenomic assembly suggests adaptation to the marine subcuticular habitat, including acquisition of Na⁺-dependent energy metabolism and loss of chemotaxis-related genes [15]. In brittle stars, an isolated *Endozoicomonadaceae* SCB has a genome characterized by genes encoding secretion systems, and the bacterium experimentally suppresses the growth of co-cultured bacteria, suggesting a role in microbial interactions within the subcuticular niche [14]. Together, these studies show that genome-resolved analyses of SCB can reveal functional traits associated with metabolic potential, niche adaptation, and possible interactions with the host or other microbes. These limited examples indicate that SCB may comprise phylogenetically diverse bacterial lineages with distinct strategies for persistence in echinoderm tissues.

In this study, we report the discovery of a novel SCB lineage associated with deep-sea holothurians of the genus *Scotoplanes*. We recovered two high-quality genomes of the dominant bacterium inhabiting the body wall of *Scotoplanes*. Based on the marked distinctiveness of its 16S rRNA gene and genome sequences, we designate this bacterium “*Candidatus* Abyssoplasma scotoplanesicola” gen. nov., sp. nov. within the phylum *Mycoplasmatota*. Microbiome profiling of different body regions showed that this bacterium is distributed throughout the body wall of *Scotoplanes* and was detected in specimens collected from multiple sites in the Northwest Pacific. Fluorescence in situ hybridization (FISH) analysis confirmed that this bacterium aggregates within host tissue on the epidermal side of the body wall, supporting its identification as a tissue-associated SCB. Comparative genomic analyses further revealed an expanded repertoire of acidic cell-surface-associated proteins in highly reduced genomes and localized genome plasticity near defense islands, providing insights into potential mechanisms of host-interface diversification.

## Materials and Methods

### Sample collection

Three specimens of *Scotoplanes theeli* Ohshima, 1915 (H4, H5, and H8) collected off Otsuchi, Iwate, Japan were previously reported [19]. Five additional *Scotoplanes* specimens (*S. theeli*, *Scotoplanes hanseni* Gebruk, 1983, and *Scotoplanes kurilensis* Gebruk, 1983) were collected using a 3-or 4-m-wide beam trawl during cruises of the R/V Shinsei Maru KS-20-15 and R/V Hakuho Maru KH-23-5 (Table 2). Specimens were identified to species based on ossicle morphology.

**Table 1.**
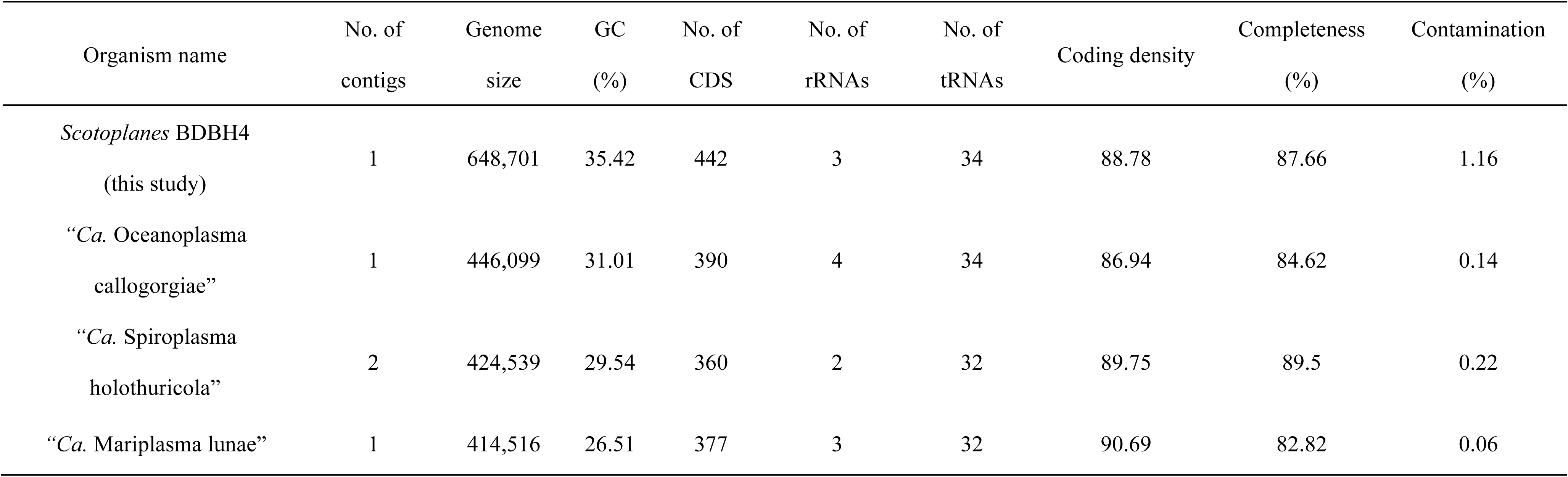
Summary of genome statistics for *Scotoplanes* BDBH4 and its closest known relative.

| Organism name | No. of<br>contigs | Genome<br>size | GC<br>(%) | No. of<br>CDS | No. of<br>rRNAs | No. of<br>tRNAs | Coding density | Completeness<br>(%) | Contamination<br>(%) |
| --- | --- | --- | --- | --- | --- | --- | --- | --- | --- |
| *Scotoplanes* BDBH4<br>(this study) | 1 | 648,701 | 35.42 | 442 | 3 | 34 | 88.78 | 87.66 | 1.16 |
| “*Ca. Oceanoplasma*<br>*callogorgiae*” | 1 | 446,099 | 31.01 | 390 | 4 | 34 | 86.94 | 84.62 | 0.14 |
| “*Ca. Spiroplasma*<br>*holothuricola*” | 2 | 424,539 | 29.54 | 360 | 2 | 32 | 89.75 | 89.5 | 0.22 |
| “*Ca. Mariplasma lunae*” | 1 | 414,516 | 26.51 | 377 | 3 | 32 | 90.69 | 82.82 | 0.06 |

**Table 2.** Sampling information for *Scotoplanes* specimens analyzed in this study with specimen ID, depth range, station name, coordinates, collection date, species identification, and whether each specimen was used for 16S rRNA gene amplicon analysis and/or fluorescence in situ hybridization (FISH).

| Cruises | Specimen<br>ID | Species | Depth range<br>(m) | Station | Coordinates, on<br>bottom | Coordinates, off<br>bottom | Date | Amplicon<br>analysis | FISH |
| --- | --- | --- | --- | --- | --- | --- | --- | --- | --- |
| R/V Shinsei Maru KS-20-15 | St1 | *Scotoplanes*<br>*theeli* | 1,594–1,590 | OT1550-1 | 39°21.02'N,<br>142°43.13'E | 39°20.11'N,<br>142°43.01'E | 1-Oct-20 | Yes | No |
| R/V Shinsei Maru KS-20-15 | St2 | *Scotoplanes*<br>*theeli* | 1,594–1,590 | OT1550-1 | 39°21.02'N,<br>142°43.13'E | 39°20.11'N,<br>142°43.01'E | 1-Oct-20 | Yes | No |
| R/V Hakuho Maru<br>KH-23-5 | Sk | *Scotoplanes*<br>*kurilensis* | 3,943–3,934 | C7 | 42°12.080'N,<br>145°40.725'E | 42°11.456'N,<br>145°40.909'E | 19-Sep-23 | Yes | No |
| R/V Hakuho Maru<br>KH-23-5 | Sh1 | *Scotoplanes*<br>*hanseni* | 5,576–5,584 | A9 | 41°02.076'N,<br>145°47.142'E | 41°01.921'N,<br>145°46.385'E | 18-Sep-23 | Yes | Yes |
| R/V Hakuho Maru<br>KH-23-5 | Sh2 | *Scotoplanes*<br>*hanseni* | 5,576–5,584 | A9 | 41°02.076'N,<br>145°47.142'E | 41°01.921'N,<br>145°46.385'E | 18-Sep-23 | Yes | Yes |

### Sanger sequencing of 16S rRNA genes

Total DNA was extracted from all eight specimens using a DNeasy Blood and Tissue Kit (QIAGEN, Germany). The three previously reported specimens were prepared as described previously [19], whereas the other five newly collected specimens were fixed on board in 99% ethanol and stored at 4 °C until DNA extraction. Near-full-length bacterial 16S rRNA gene fragments were amplified using primers 27F/1492R [21] (Table S1). Purified PCR products were Sanger sequenced using primers 533F/907R [22] for the previously reported specimens. In contrast, newly designed “*Ca*. A. scotoplanesicola”-targeted primers were used for the additional specimens. Further details are provided in the Supplementary Materials and Methods.

### Metagenome analysis and genome reconstruction

Total DNA of the body walls of *S. theeli* specimens H4, H5, and H8 was extracted using a Genomic-tip 500/G (QIAGEN, Germany). Paired-end libraries were prepared and sequenced on an Illumina MiSeq platform (2 x 301 bp). Reads from each sample were assembled de novo using MetaPlatanus v1.0.4 [23], and contigs >1kb were binned using MetaBAT2 v2.12.1 [24]. One *Mycoplasmatota* bin was identified from each sample using GTDB-tk2 v2.1.1 [25] (Table S3), and no additional bacterial bins were recovered. Bin quality was assessed using CheckM2 v1.1.0 [26], and the corresponding 16S rRNA gene sequences from H4, H5, and H8 were identified in the bins using BLASTn.

To reconstruct a complete circular genome, sample H4 was sequenced on a HiSeq X Ten platform (2 x 151 bp) using 3-and 6-kb mate-pair libraries. Reads were assembled using Platanus_B v1.3.2 [27], yielding a single 643.6-kb scaffold with gaps and an identical 16S rRNA gene sequence. Read mapping of the paired-end and mate-pair libraries supported scaffold circularity, and gaps were manually curated based on mapped read-pair distances and orientations. This yielded a complete, gap-free circular genome sequence of the body-wall-dominant bacterium from the H4 specimen (BDBH4). Protein-coding sequences (CDSs) were predicted by using PROKKA v1.12 [28]. rRNA genes were identified using Barrnap and Infernal [29,30].

### Phylogenetic analysis

Representative 16S rRNA sequences for each family within *Mycoplasmatota,* together with two within *Bacillota* as outgroup sequences, were obtained from NCBI. These sequences and the three sequences from H4, H5, and H8 were aligned using MAFFT v7.526 [31]. A maximum-likelihood (ML) 16S rRNA phylogeny was reconstructed using IQ-TREE v2.3.6 [32] with 1,000 bootstrap replicates and the best-fit model, GTR+F+I+R4, selected by ModelFinder [33].

For phylogenomic analysis, 47 *Mycoplasmatota* genomes were selected from the 159 genomes listed in Table S2. Orthologous groups (OGs) in these genomes and BDBH4 were inferred using OrthoFinder v2.5.5 with default parameters [34]. Amino acid sequences of 56 OGs conserved across all the genomes were individually aligned with MAFFT v7.526 (--auto) [31], trimmed using trimAl v1.4 with the-nogaps option [35], and concatenated. An ML phylogeny was reconstructed using RAxML-NG v0.9.0 under the LG+I+G4m model selected by automatic model selection [36]. Nodal support was assessed with 1,000 bootstrap replicates (--bs-trees 1000).

### Gene annotation

Gene annotation of BDBH4 (Tables S4–S9) and BDBH8 were performed as follows. COGs were assigned using COGclassifier [37]. KEGG KO was assigned using eggNOG-mapper v2 and KOfam [38,39]. Pfam domains were identified using hmmscan [40]. The defense system was identified using DefenseFinder [41]. CRISPR spacers and Cas proteins were searched using the CRISPRcasFinder online [42]. Inverted repeats were identified using EMBOSS einverted [43]. Subcellular localization and membrane topology of encoded proteins were predicted using DeepTMHMM v1.0.24 [44], SignalP v6.0h [45], and Phobius v1.01 [46] with default parameters.

For comparative genomics, 159 *Mycoplasmatales* genomes were collected from NCBI. PROKKA v1.1 [28] and eggNOG-mapper v2 [39] were used for CDS detection and gene annotation. OGs were identified among the 159 genomes and BDBH4 using OrthoFinder v2.5.5 with default parameters [34].

### Structural analysis

Protein structures were predicted using AlphaFold v3.0.2 [47] with customized alignments. Structural modules were defined based on predicted aligned error (PAE), and segment similarities were evaluated using Foldseek v10.941cd33 [48], as described in the Supplementary Materials and Methods.

### 16S rRNA amplicon sequencing

*Scotoplanes* specimens were dissected to collect tissues from different body parts. Tissue samples were lyophilized, homogenized, and subjected to DNA extraction using either the MPure 12 system with an MPure Bacterial DNA Extraction Kit (MP Biomedicals, USA) or a Lab-Aid 824s DNA Extraction Kit (Zeesan Biotech, China), depending on the host specimen. Amplicon libraries targeting the V3–V4 region of the 16S rRNA gene were prepared by two-step tailed PCR using the primer sets listed in Table S1. The obtained reads were processed using DADA2 [49] and analyzed using QIIME 2 [50].

### Hybridization chain reaction-fluorescence in situ hybridization (HCR-FISH)

Body-wall and gonadal tissues were excised from three specimens (Sk, Sh1, and Sh2; Table 2), fixed on board in 4% paraformaldehyde phosphate buffer solution at 4°C for 8–12 h, and stored in 70% ethanol at 4°C until further processing. A new 16S rRNA-targeted probe was designed for “*Ca.* A. scotoplanesicola”. Fixed tissues were decalcified, paraffin-embedded, serially sectioned at 10 µm, and mounted on slides. FISH was performed using the designed probe, the universal bacterial probe EUB338, and the negative-control probe NonEUB338 [51,52]. After hybridization and signal amplification, sections were mounted with antifade reagent and observed using an Olympus FV3000 confocal microscope in super-resolution mode.

## Results

### Detection of a body-wall dominant bacterial lineage from *Scotoplanes theeli*

We explored the body-wall bacterial community of *S. theeli* by PCR amplification and Sanger sequencing targeting the full-length 16S rRNA gene from three *S. theeli* specimens (H4, H5, and H8) collected off Otsuchi [19]. PCR amplifications yielded a single band with an expected length from all three specimens. Sequencing of the PCR products produced unambiguous signals across the full length, and among the obtained sequences, H4 and H8 were identical, while H5 differed from both by two nucleotides (Fig. S1). These results indicated the dominance of a single bacterial lineage in the body-wall microbiome of the three specimens, hereafter referred to as the *Scotoplanes* body-wall-dominant bacterium, abbreviated as *Scotoplanes* BDB.

BLAST searches of the 16S rRNA gene sequences of the *Scotoplanes* BDB against the NCBI nt database indicated that this bacterial lineage belongs to the phylum *Mycoplasmatota*. The closest known relative was “*Candidatus* Oceanoplasma callogorgiae”, originally discovered in the mesoglea of deep-sea corals [53], with 82.3% sequence identity to *Scotoplanes* BDB. A maximum-likelihood phylogenetic tree based on full-and near-full-length 16S rRNA gene sequences placed the three *Scotoplanes* BDB sequences within the order *Mycoplasmatales* (Fig. 1A). Marine animal–associated species in this order have been reported from the Hominis group, the Pneumoniae group, and “*Ca.* Oceanoplasmataceae” [53]. The three *Scotoplanes* BDB sequences clustered within “*Ca.* Oceanoplasmataceae”, a family that includes species associated with jellyfish (“*Ca.* Mariplasma lunae”) [54], deep-sea corals (“*Ca.* Oceanoplasma callogorgiae”) [53], crown-of-thorns starfish [55], and sea cucumbers (“*Ca.* Spiroplasma holothuricola”) [56].

**Figure 1.**
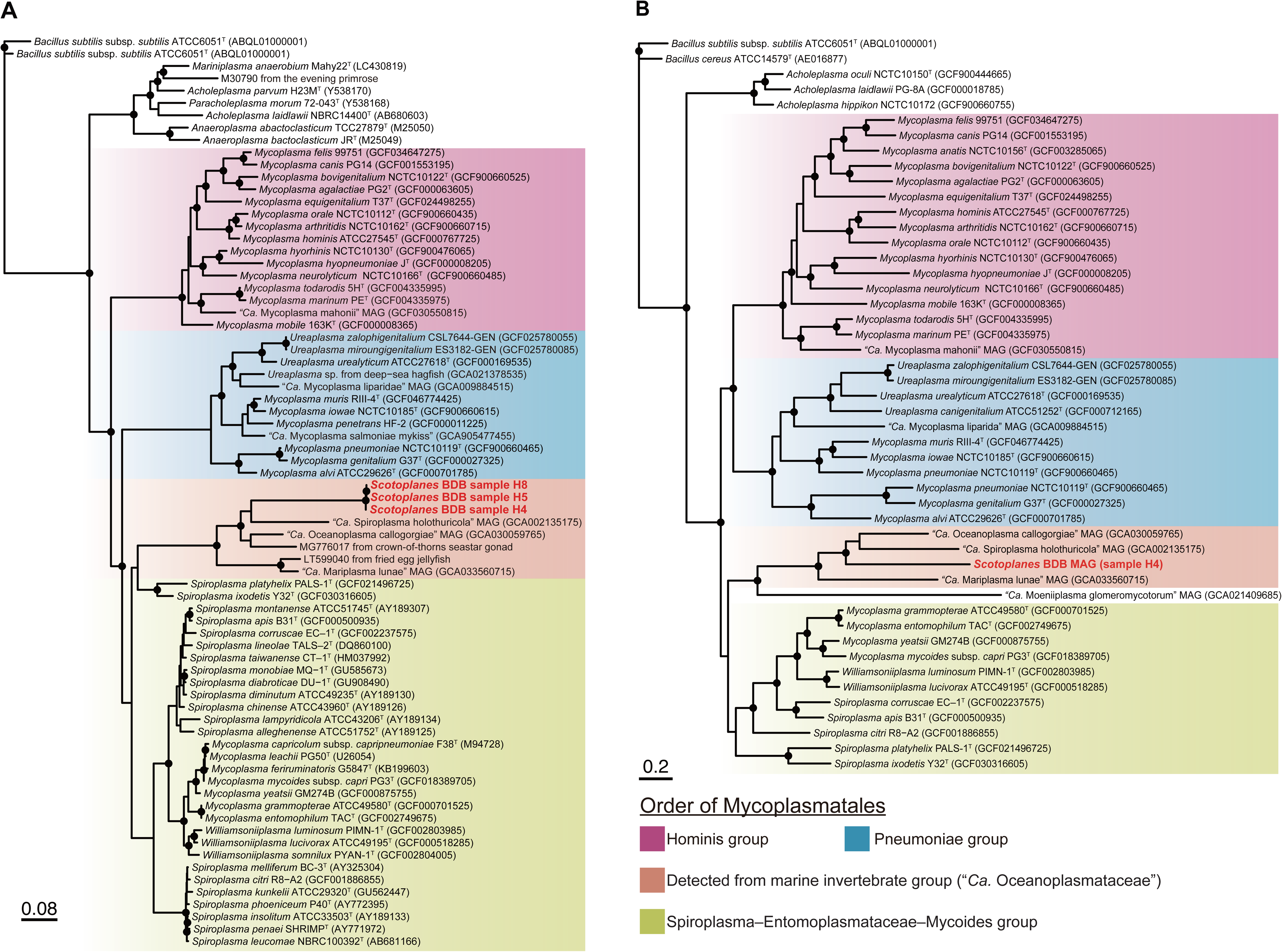
Maximum-likelihood phylogenies of the phylum *Mycoplasmatota*. Phylogenies were inferred from (A) 16S rRNA gene sequences and (B) concatenated amino acid sequences of 56 conserved single-copy genes. Colored backgrounds indicate major groups within the order *Mycoplasmatales*. Filled circles denote nodes with ultrafast bootstrap (UFBoot2) support values >95%. Accession numbers are shown in parentheses. *Scotoplanes* BDB is highlighted in bold red.

### Reconstruction of the complete circular genome of *Scotoplanes* BDB

To gain genomic insights into the *Scotoplanes* BDB, we assembled metagenomic sequence reads from samples H4, H5, and H8 and obtained three *Mycoplasmatales* genomic bins (Table S3). The bin from H4 showed the highest completeness (87.13%) and assembly continuity, comprising two contigs of 466 kb and 157 kb. To close the assembly, additional 3-kb and 6-kb mate-pair libraries were generated for sample H4. Using both paired-end and mate-pair reads, we obtained a complete circular genome assembly of the *Scotoplanes* BDB. The genome, hereafter referred to as the BDBH4 genome, was 648,701 bp in size, had a GC content of 35.42%, and was predicted to encode 442 CDSs.

Phylogenomic analysis based on 56 conserved single-copy genes showed that BDBH4 formed a well-supported clade with members of “*Ca*. Oceanoplasmataceae” but was clearly separated from the three available genomes in the family (Fig. 1B). The average nucleotide identity (ANI) between the BDBH4 genome and the three known “*Ca.* Oceanoplasmataceae” genomes ranged from 62.9% to 63.6%, indicating substantial genomic divergence from the other members [57]. Together with its phylogenetic placement, this divergence supports the designation of *Scotoplanes* BDB as a novel genus-level lineage within “*Ca.* Oceanoplasmataceae”. We therefore propose the new genus and species “*Candidatus* Abyssoplasma scotoplanesicola” gen. nov., sp. nov. “*Abyssoplasma*” combines a reference to the dark deep sea, from which the symbiont was recovered, with “-plasma”, reflecting its phylogenetic affiliation with *Mycoplasmatales*. The species epithet “*scotoplanesicola*” denotes an organism inhabiting *Scotoplanes*.

### Comparative genomics and metabolic potential of “*Ca*. Abyssoplasma scotoplanesicola”

We compared the genomic features of BDBH4 with those of related lineages. Based on comparison with 159 genomes, the BDBH4 genome was smaller and contained fewer CDSs than most genomes in the order *Mycoplasmatales* (Fig. 2A). However, in comparison with more closely related genomes from the family “*Ca.* Oceanoplasmataceae”, the BDBH4 genome was approximately 200 kb larger, encoded more CDSs, and had a higher GC content (Table 1). The genome contained three rRNA genes (5S, 16S, and 23S) that were not organized in a canonical operon but were dispersed across the genome, separated by intervals of 43–117 kb (Figs. S3, S4). This separation may reflect past genome rearrangement, as suggested by a pair of inverted repeats identified near the 16S rRNA gene (Fig. S4), although no integrase or recombinase genes were identified in the genome. The genome contained 34 tRNA genes covering all 20 amino acids. The genomic coding density, estimated completeness, and estimated contamination were comparable to those of other members of the “*Ca.* Oceanoplasmataceae” (Table 1).

**Figure 2.**
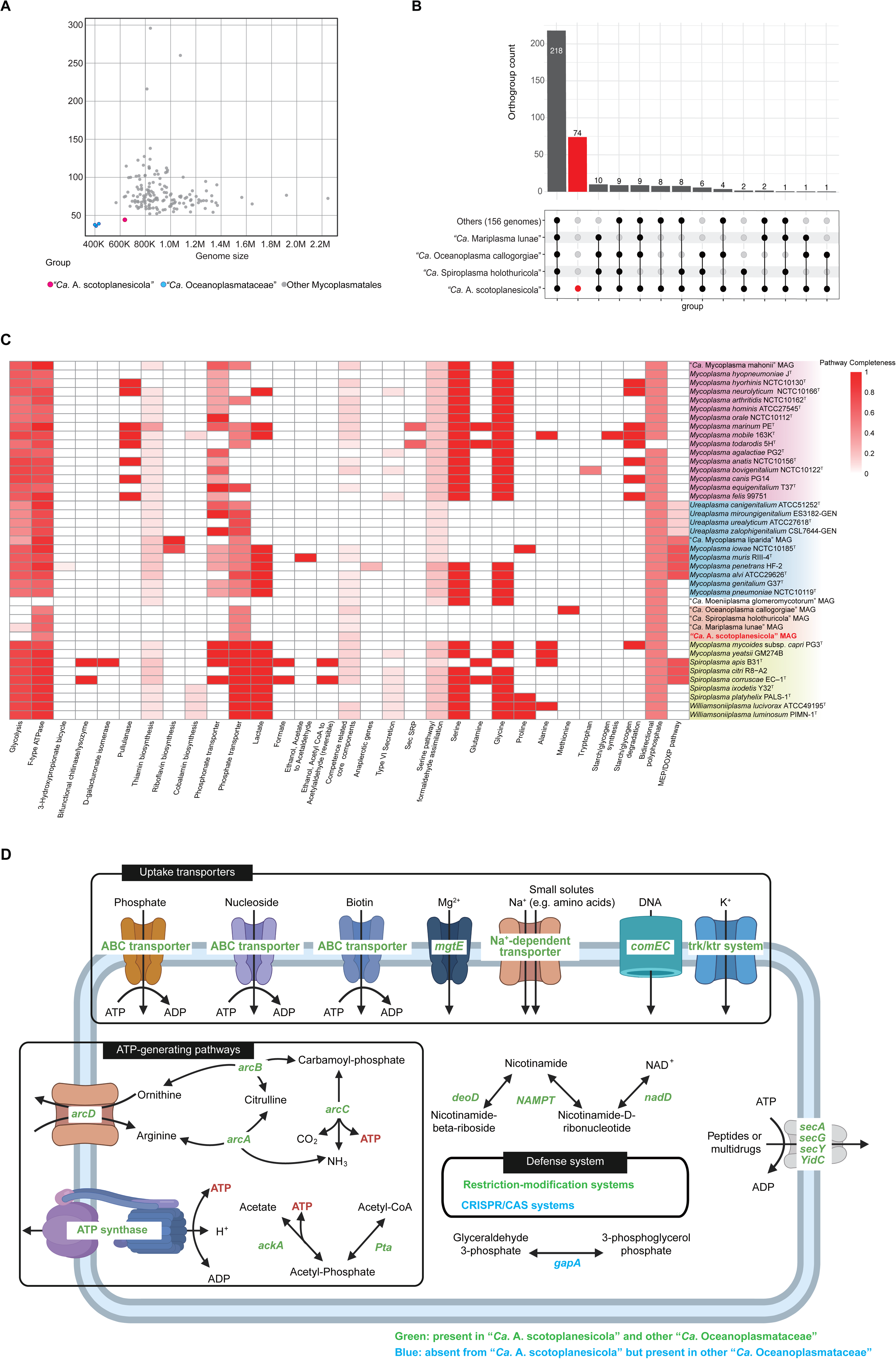
Genome features and predicted metabolic capabilities of “*Ca*. A. scotoplanesicola”. (A) Genome size and CDS number of “*Ca.* A. scotoplanesicola” compared with representative *Mycoplasmatales* genomes. (B) UpSet plot showing orthologous group (OG) sharing patterns between the BDBH4 genome, three related marine invertebrate-associated group genomes, and 156 other *Mycoplasmatales* genomes (Others). Vertical bars indicate the number of OGs in each intersection, with values shown above the bars. The dot-and-line matrix below the bar plot indicates the genomes included in each intersection. The red bar indicates “*Ca*. A. scotoplanesicola”-specific OGs, including an OG containing long-chain genes. (C) Metabolic and functional pathway completeness across *Mycoplasmatales* genomes, showing a heatmap for selected metabolic, biosynthetic, and cellular pathways. The completeness is estimated using KEGG-Decoder. Values represent the fraction of required KEGG orthologs detected for each pathway (0 = not detected, 1 = complete); color intensity increases with completeness. Genome labels are colored according to the major *Mycoplasmatales* groups. Genomes are ordered following the phylogenomic relationships shown in Fig. 1. (D) Schematic representation of predicted metabolic and cellular functions in “Ca. A. scotoplanesicola”, highlighting nutrient uptake systems, the arginine deiminase pathway, acetate production via *Pta/AckA*, ATP synthase, NAD salvage, DNA uptake, secretion-related genes, and defense-related genes. Green gene labels indicate genes present in “*Ca*. A. scotoplanesicola” and other Oceanoplasmataceae, whereas blue labels indicate genes absent from “*Ca*. A. scotoplanesicola” but present in other “*Ca.* Oceanoplasmataceae”. Figure partially created in BioRender. Y. Y. (2026). https://BioRender.com/q77k751

To characterize the metabolic capabilities of the BDBH4 genome, we performed homology-based gene annotation using multiple databases. We then clustered genes from “*Ca*. A. scotoplanesicola” and the 159 *Mycoplasmatales* genomes into orthologous groups (OGs). The BDBH4 genome contained 353 OGs, of which 74 (21.0%) were not identified in any of the other analyzed genomes (Fig. 2B). This indicates that the BDBH4 genome contains a substantial set of lineage-specific genes that are absent even from the other *Mycoplasmatales* genomes.

The BDBH4 genome exhibited a highly reduced metabolic repertoire relative to the other *Mycoplasmatales* genomes, while sharing a broadly similar set of metabolic orthologs with known “*Ca.* Oceanoplasmataceae” genomes (Fig. 2C). Genes encoding central carbon metabolism and carbohydrate-based metabolic pathways for biomass or energy production, including glycolysis, the TCA cycle, and carbohydrate fermentation pathways, were not detected. In contrast, the genome retained genes for the arginine deiminase (ADI) pathway, F-type ATP synthase, and *ackA* (Fig. 2D). The ADI pathway genes included the complete set for arginine-to-ornithine conversion (*arcABCD*), suggesting a potential carbohydrate-independent ATP-generating route through arginine catabolism [58–60]. Genes encoding all F-type ATP synthase subunits, including a putative ε subunit (*atpC*), were also detected. In non-fermentative *Mycoplasmatota* such as *Ureaplasma*, F-type ATP synthase can be coupled to ion gradients generated by ammonia/ammonium metabolism [61]. The retained *ackA* encodes acetate kinase and suggests a potential capacity for substrate-level phosphorylation, although its metabolic context remains unclear.

Genes for de novo amino acid biosynthesis were largely absent, whereas *metK*, encoding S-adenosylmethionine (SAM) synthetase, was retained, suggesting a capacity for SAM production. No complete pathways for vitamin, cofactor, or coenzyme biosynthesis were detected, although NAD precursor salvage/activation and conversion of riboflavin to FMN and FAD were apparent (Fig. 2C). Consistent with this reduced biosynthetic capacity, the BDBH4 genome retained 12 transporter genes, including ABC transporters for nucleotides, biotin, and phosphate (*nupABC, bmpA*), ion transporters (*mgtE, Ktr*), and four copies of sodium ion-dependent transporters (SNF), suggesting potential uptake capacities for nutrients, ions, and other substrates (Tables S5-7, Fig. 2D). The genome also encoded *comEC*, a core component of bacterial DNA uptake systems. As defense-related features, multiple restriction–modification (RM) systems were identified in the genome.

The overall pattern of metabolic gene content was broadly shared among the “*Ca.* Oceanoplasmataceae” genomes. However, the BDBH4 genome lacked an ortholog of glyceraldehyde-3-phosphate dehydrogenase (*gapA*), which was conserved in 151 of the 159 *Mycoplasmatales* genomes analyzed. BDBH8, a genomic bin recovered from sample H8, consisted of three long contigs and had high estimated completeness (87.13%). BDBH8 shared the same metabolic OG profile as BDBH4, including the absence of *gapA*. This suggests that the absence of *gapA* is a lineage-specific feature of “Ca. A. scotoplanesicola”. In addition, although CRISPR-Cas systems were conserved in the other analyzed genomes of “*Ca.* Oceanoplasmataceae”, these systems were not identified in either BDBH4 or BDBH8.

### Expanded acidic cell-surface-associated protein groups and genome plasticity in “*Ca*. A. scotoplanesicola”

The average CDS length of BDBH4 was 433.3 amino acids (aa), exceeding the maximum value observed among the other “*Ca.* Oceanoplasmataceae” genomes (351.8 aa; Fig. S2A). This elevated average CDS length was attributable primarily to the presence of several large CDSs (Figs. S3, S4). BDBH4 contained 13 CDSs longer than 2,000 aa, whereas the other “*Ca.* Oceanoplasmataceae” genomes contained only one such CDS in total. By contrast, 518 CDSs longer than 2,000 aa were found across the other *Mycoplasmatale*s genomes (Fig. S2B), suggesting that large CDSs are not unusual across *Mycoplasmatales*. The 13 large CDSs in BDBH4 were predicted to encode transmembrane proteins but lacked specific functional annotations.

We further examined functionally unannotated membrane-associated proteins and their OGs in BDBH4. We identified six nonsingleton OGs of membrane-associated proteins, comprising 37 sequences >200 aa in total. These groups, designated A–F, were tentatively categorized as acidic cell-surface-associated proteins (ASAPs) based on their predicted low isoelectric points (pIs; ranging from 3.92 to 5.34, excluding a group E outlier) and large fractions predicted to be extracellular (ranging from 74.6% to 98.8%; Figs. 3A, S5). In particular, groups A, B, and C included large proteins of up to 5,476, 2,819, and 2,387 aa, respectively, whereas proteins in group E were much smaller and more uniform in size, with the largest being 341 aa. Together, these groups accounted for a substantial fraction of the genome, representing up to 27.6% of the total genome size. Based on their low pI values, these proteins are expected to be net negatively charged under near-neutral conditions. All proteins in groups A, B, C, and E were predicted to have N-terminal signal peptides (SPs) for secretion and C-terminal transmembrane regions (TMs), and these groups were identified as BDBH4-specific OGs. In groups D and F, most proteins lacked C-terminal TMs but contained either an N-terminal TM or an N-terminal SP (Figs. 3A, S5). An ortholog of group D was identified in “*Ca.* Oceanoplasma callogorgiae” and two orthologs of group F were identified in “*Ca.* Spiroplasma holothuricola”. Both species belong to “*Ca.* Oceanoplasmataceae”. ASAPs lacked functional annotations, except for two group F proteins that were partially annotated.

**Figure 3.**
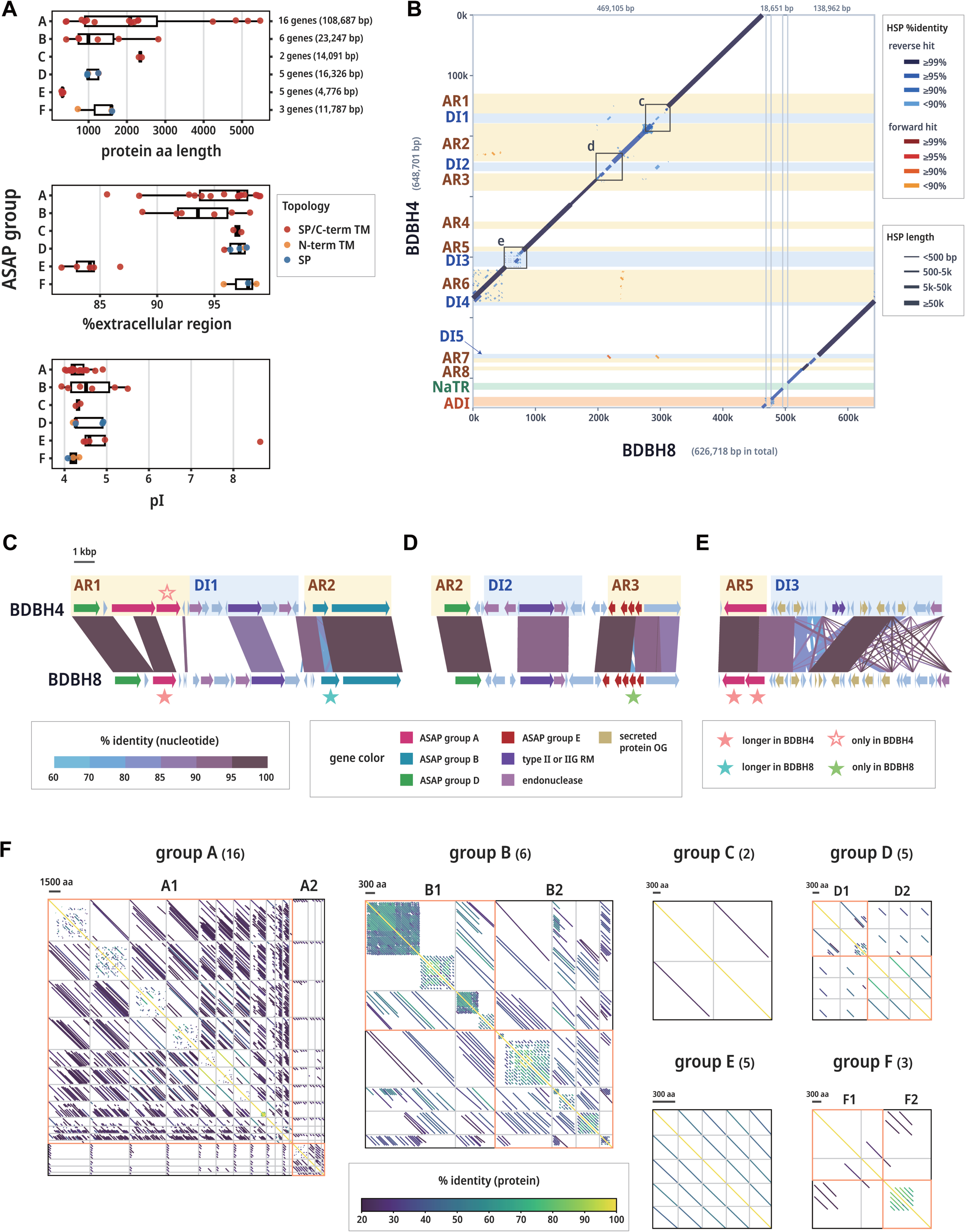
Sequence properties of ASAPs and comparison between two “*Ca.* A. scotoplanesicola” genomes. (A) Protein length, predicted extracellular-region fraction, and pI of ASAPs summarized by group. Each point represents an individual protein >200 aa and is colored according to its predicted topology: proteins with both signal peptide-like and C-terminal transmembrane regions, proteins with only an N-terminal transmembrane region, and proteins with only a signal peptide-like region (SP). The number of genes and total base pairs are shown on the right side of the upper panel. (B) BDBH8-to-BDBH4 nucleotide similarity visualized as a BLASTn dotplot. BLASTn was run with DUST filtering enabled. Contigs were separated by fixed visual gaps to distinguish individual BDBH8 contigs. Only HSPs of at least 200 bp were plotted. BDBH4 genomic regions corresponding to ASAP regions (ARs), defense islands (DIs), NaTR, and ADI are highlighted. Detailed views of the BDBH4 genome are shown in Fig. S4. (C–E) Expanded maps of three localized genome rearrangement sites between BDBH4 and BDBH8, corresponding to the regions indicated in panel (B). HSPs within each region are shown. Genes encoding ASAPs, type II or IIG RM systems, endonucleases, and members of a secreted-protein OG are highlighted. Corresponding ASAP pairs were assigned based on nucleotide similarity and genomic collinearity. ASAPs present only in one genome or differing in length by >100 aa between genomes are indicated with stars. (F) Dotplots of ASAP groups generated from within-group all-vs-all BLASTp searches using proteins >200 aa. Member proteins were concatenated on both axes, and BLASTp HSPs were drawn according to their query and subject coordinates. Inter-protein hits were visualized from standard gapped BLASTp searches, whereas self-comparisons were visualized from separate ungapped BLASTp searches (-ungapped-word_size 3) to emphasize internal repeat signals. BLASTp was run with SEG masking and composition-based statistics disabled (-seg no-comp_based_stats F), and only hits with E-value <1e-3 were plotted.

Genes encoding these protein groups were clustered in eight genomic regions, hereafter referred to as ASAP regions (ARs; AR1–AR8; Fig. S4). In addition, we identified five defense islands (DIs; DI1–DI5; Fig. S4), each encoding genes for type II or IIG RM systems and endonucleases. Notably, five of the eight ARs (AR1–AR3, AR5, AR7) were located adjacent to DIs.

We compared the overall genome structures of BDBH4 and BDBH8 by mapping BDBH8 contigs onto the BDBH4 genome using BLASTn (Fig. 3B). The two genomes showed high overall similarity (ANI = 99.0%) and identical 16S rRNA gene sequences. However, four DIs (DI1–DI3, DI5) and their neighboring regions, including DI-adjacent portions of ARs (AR1–AR3, AR5), showed substantial divergence between the two genomes (Figs. 3B–E, S4). In DI5, a gene encoding a type IIG RM system was absent from BDBH8. Together, these patterns suggest that DIs and adjacent ARs constitute hotspots of localized genome rearrangement in “*Ca.* A. scotoplanesicola”.

To further inspect ASAPs, homologous regions within each group were identified using BLASTp. Visualization of the positions of these homologous regions suggested that groups A and B, as well as some members of groups D and F, contain repeat units (Fig. 3F). This pattern suggests that some ASAPs have modular architectures, potentially contributing to their large size and sequence variability.

To examine ASAPs beyond sequence-level homology, protein structures were predicted using AlphaFold3. The predicted structures were generally supported by moderate to high confidence scores (with mean pLDDT values ranging from 53.1 to 85.5; Table S10), except for those of group C (mean pLDDT < 40). We therefore excluded the predicted structures of group C from subsequent analyses. Based on the predicted structures, we defined 3D modular segments, ranging from 70 to 292 aa, which were supported by low mean PAE values within each segment (Figs. 4A–B). Structural similarity among the segments was evaluated using Foldseek TM-score (Fig. 4E). Clustering based on the similarity yielded three major clusters, hereafter referred to as clusters α, β, and γ, comprising 281, 44, and 26 segments, respectively. Segments belonging to the largest cluster, cluster α, were identified in groups A, B, D, and E, indicating repeated use of similar structural units in proteins of these groups.

**Figure 4.**
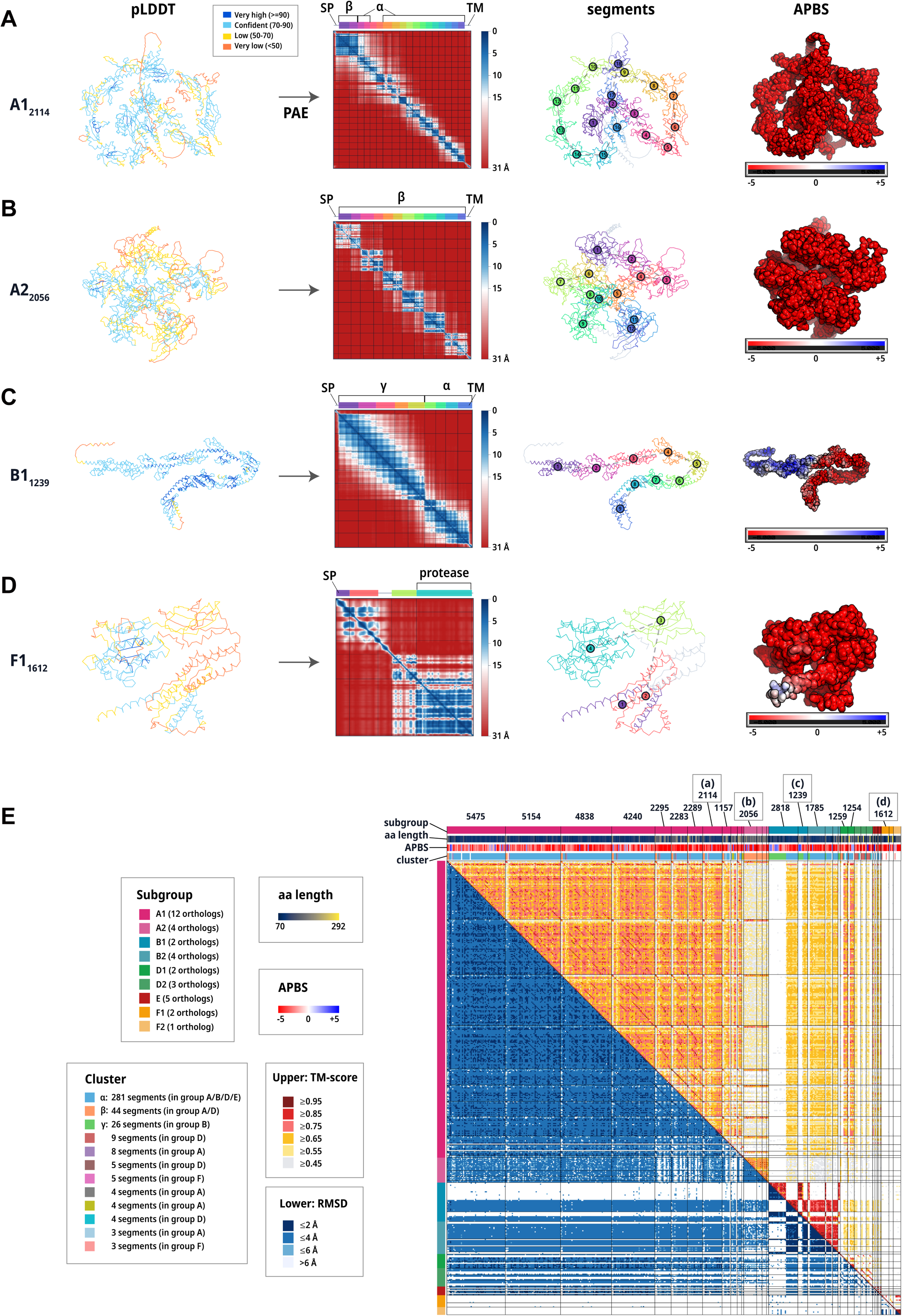
Structural insights into ASAPs. (A–D) Representative AlphaFold3-predicted structures and structural features of ASAPs. From left to right, each row shows the predicted structure colored by pLDDT score, the PAE matrix with PAE-based modular segmentation, the structure colored by PAE-defined modular segment, and the electrostatic potential calculated using APBS. Representative proteins are shown for (A) a 2,114-aa protein in subgroup A1, (B) a 2,056-aa protein in subgroup A2, (C) a 1,239-aa protein in subgroup B1, and (D) a 1,612-aa protein in subgroup F1. Colored bars above the PAE matrices indicate the identified modular segments. Clusters of modular segments in panel (E) and the segment containing the protease domain are shown above the colored bars. Predicted signal peptide-like regions (SPs) and transmembrane regions (TMs) are also shown above the colored bars. Electrostatic potentials are shown on a fixed scale from −5 to +5 kT/e. (E) Heatmap showing all-vs-all structural comparisons among 419 PAE-defined modular segments from 35 ASAP proteins >200 aa, excluding group C. The upper triangle of the heatmap shows TM-scores, and the lower triangle shows RMSD values. Bars above the heatmap indicate subgroup, amino acid length, electrostatic potential, and structural cluster for each segment. Amino acid lengths are indicated for proteins >1,000 aa. The proteins shown in panels (A)–(D) are also indicated.

Group A proteins were classified into two subgroups. In subgroup A1, one or two N-terminal segments belonged to cluster β, whereas most remaining segments belonged to cluster α (Fig. 4A). In contrast, subgroup A2 consisted entirely of cluster β segments (Fig. 4B). Group B proteins were also classified into two subgroups based on the presence (B1) or absence (B2) of a characteristic positively charged N-terminal region (Fig. S5). In subgroup B1, this region corresponded to N-terminal segments enriched in cluster γ, whereas most remaining segments belonged to cluster α (Fig. 4C). Among the three group F proteins, two sequences, designated subgroup F1, contained a serine protease domain (Fig. 4D). Although their N-terminal regions were predicted either as SPs or as N-terminal TMs, depending on the prediction software used, the serine protease domain was located in the C-terminal region, suggesting that the active site is likely exposed extracellularly regardless of the predicted N-terminal topology.

### Distribution of “*Ca.* A. scotoplanesicola” across body parts of *Scotoplanes* holothurians sampled from the Northwest Pacific

Bacterial community compositions in different body parts of *Scotoplanes* spp. were analyzed using five specimens, including two *S. theeli*, one *S. kurilensis*, and two *S. hanseni* specimens (Table 2), collected from three sampling sites in the Northwest Pacific Ocean (Fig. 5A). Seven tissue types were sampled from *S. theeli*: body wall, papilla, tube foot, foregut, midgut, hindgut, and gonad. Only body wall samples were collected from *S. kurilensis* and *S. hanseni* (Fig. 5B). All 17 tissue samples were subjected to microbial community analysis targeting the V3–V4 region of the 16S rRNA gene. The gonadal samples of *S. theeli* were excluded from subsequent data analyses because of insufficient sequencing quality. After quality filtering, 504,106 reads were retained in total (Table S11).

**Figure 5.**
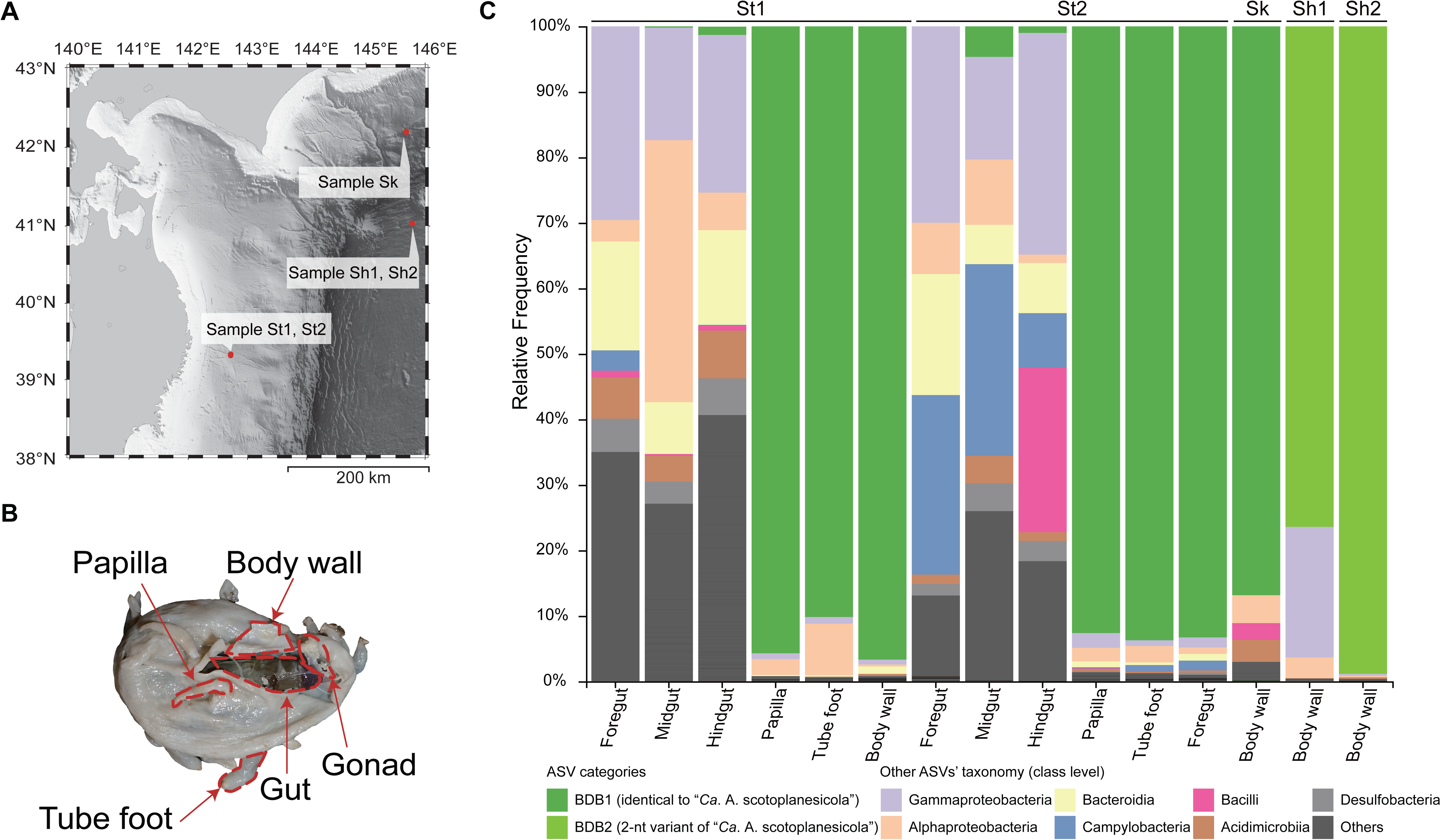
Tissue sampling and amplicon profiles of “*Ca*. A. scotoplanesicola”. (A) Map of sampling stations; letters (St1, St2, Sk, Sh1, Sh2) correspond to individual specimens/sites used in this study. (B) Dissected specimen of *Scotoplanes* spp. showing the tissues used for 16S rRNA gene amplicon sequencing (papilla, tube foot, body wall, gut, and gonad). (C) Stacked bar plots showing class-level taxonomic composition (relative frequency) of amplicon sequence variants (ASVs) across tissues from each specimen. Green indicates the BDB1, and light green indicates a closely related BDB2 differing by two nucleotides; all other ASVs are grouped by bacterial class, with low-abundance taxa combined as “Others.”

Across all five specimens, a single amplicon sequence variant (ASV) identical to or closely related to “*Ca.* A. scotoplanesicola” BDBH4 was consistently dominant in the body wall (76.4–98.9%). In the body wall samples from St1, St2, and Sk, the dominant ASV (BDB1) was identical to the BDBH4 sequence, whereas the dominant ASV in Sh1 and Sh2 (BDB2) differed from it by two nucleotide positions (Fig. 5C). In *S. theeli*, the BDBH4-identical ASV also dominated the tube feet and papillae, suggesting that “*Ca*. A. scotoplanesicola” broadly colonized the external body wall. In contrast, it was detected only at low relative abundance in the midgut and hindgut (0.19–4.65%), indicating a strong spatial preference for the external body surface.

To further assess sequence conservation beyond the amplicon region, we determined near-full-length 16S rRNA gene sequences from the body-wall tissues of the five specimens. We successfully obtained 1,465 bp of the 16S rRNA gene sequences using newly designed primers (Fig. S1). These sequences showed at least 99.6% nucleotide identity to those previously obtained from specimens H4, H5, and H8, indicating that the “*Ca*. A. scotoplanesicola” populations were highly conserved across sites from off Iwate to the Kuril–Kamchatka Trench.

### Localization of “*Ca*. A. scotoplanesicola” within the body wall

To examine the localization of “*Ca*. A. scotoplanesicola” in the body wall, HCR-FISH was performed on histological sections using a “*Ca*. A. scotoplanesicola”-targeted probe together with a universal bacterial probe. HCR-FISH was conducted on all specimens, with detailed analyses focused on the best-preserved *S. hanseni* specimens (Table 2).

Signals of “*Ca*. A. scotoplanesicola” were localized on the epidermal side of the body wall and formed spherical aggregates of 2–3 µm across (Figs. 6, S6). Assuming that “*Ca*. A. scotoplanesicola” cells are comparable in size to other members of the order *Mycoplasmatales*, which generally range from 200 to 500 nm [62], each aggregate was large enough to contain multiple bacterial cells. Based on host autofluorescence, the host cuticular layer was not clearly distinguishable, but the signals were consistently observed within the host tissue, supporting tissue-associated localization rather than superficial attachment. The localization of these aggregates was comparable to that of SCB aggregates previously reported in the subcuticular layers of echinoderms [63,64]. Taken together, these observations indicate that “*Ca*. A. scotoplanesicola” represents SCB associated with *Scotoplanes* spp.

**Figure 6.**
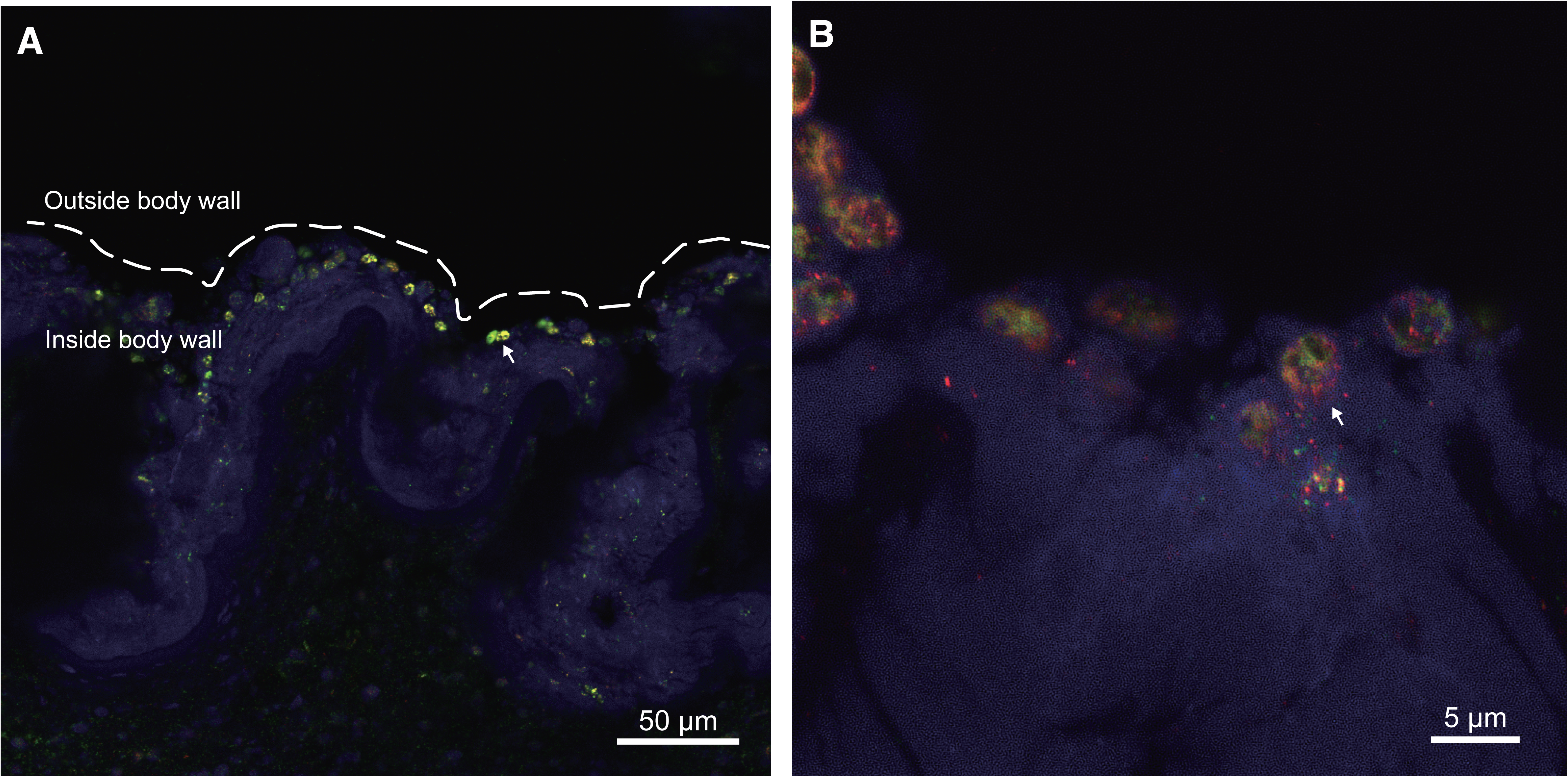
Fluorescence in situ hybridization (FISH) of the body wall of *Scotoplanes hanseni.* (A, B) Representative body-wall sections showing aggregates of “*Ca*. A. scotoplanesicola” (SCB; white arrows). Signals are shown as merged images: the universal bacterial probe EUB338 (red) and the “*Ca*. A. scotoplanesicola”–designed probe (green); blue indicates host tissue autofluorescence. White dashed lines indicate the outer boundary of the body wall, corresponding to the body surface.

We also performed HCR-FISH on gonadal samples. Although 16S rRNA gene amplification from gonadal tissue failed, “*Ca*. A. scotoplanesicola” signals were detected in the gonads (Figs. S6, S7). The signals appeared as aggregates similar to those observed in the body wall, suggesting the presence of this SCB in both the body wall and gonads.

## Discussion

This study identifies a dominant *Mycoplasmatota* symbiont inhabiting the body wall of deep-sea holothurians and provides a genome-resolved and spatially resolved characterization of holothurian-associated SCB. Body-wall samples of *Scotoplanes* spp. from the Northwest Pacific were dominated by a single bacterial species, “*Ca.* A. scotoplanesicola”, and FISH showed that this bacterium was localized on the epidermal side of the body wall. Together, these findings establish “*Ca.* A. scotoplanesicola” as a dominant SCB of *Scotoplanes* and extend genome-resolved SCB research to holothurians, a major echinoderm class in which the metabolic potential and host-tissue associations of SCB have remained poorly resolved.

“*Candidatus* Abyssoplasma scotoplanesicola” belongs to the family “*Ca.* Oceanoplasmataceae” within *Mycoplasmatota*. Although *Mycoplasmatota* are well known as pathogens of humans, livestock, and plants [65], recent studies have revealed diverse symbiotic *Mycoplasmatota* in marine invertebrates, including coastal starfish [66], a basket star [67], and intertidal brittle stars [14]. Members of “*Ca.* Oceanoplasmataceae” have been detected in diverse marine invertebrate hosts, including corals [53], jellyfish [54], crown-of-thorns starfish [55], and holothurians in this study. Despite this broad host range, available genomes of this family show highly reduced and broadly similar metabolic capabilities. This reduction and similarity suggest that these bacteria likely share a conserved host-associated metabolic strategy that enables associations with diverse marine invertebrates, rather than being tightly specialized to a single host lineage or tissue environment.

In contrast to this broadly conserved metabolic potential, the expanded repertoire of ASAPs, including members exceeding 5,000 aa, represented a distinctive feature of “*Ca.* A. scotoplanesicola”. The sequence-level repeat units and repeated structurally similar modules support modular diversification of ASAPs. In several *Mycoplasmatota* lineages, including *Mycoplasma*, *Spiroplasma*, and *Ureaplasma*, large surface-associated proteins with repetitive architectures have been reported and are involved in antigenic variation, motility, or host–cell interactions [68–70]. Acidic, repetitive surface proteins have also been described in other bacteria, such as *Staphylococcus aureus* within *Bacillota*, where they are involved in cell-surface adhesion and can exhibit calcium-dependent aggregation [71]. Since acidic proteins are likely negatively charged under near-neutral conditions, Ca²⁺ may promote electrostatic bridging among acidic surface domains and modulate interactions with bacterial cells or host surfaces. In addition, diverse types of surface protein glycosylation have been reported in mycoplasmas [72], suggesting that ASAPs may also be glycosylated. Together, these features suggest that ASAPs could form flexible, adhesive surface layers that contribute to bacterial aggregation or interactions with host tissues.

The close genomic proximity between defense islands (DIs) and ASAP regions (ARs) suggests a possible evolutionary link between DI plasticity and ASAP diversification. DIs are often associated with mobilome components [73] and represent dynamic genomic loci enriched in defense functions against mobile genetic elements (MGEs) [74]. Moreover, some defense systems, particularly RM systems, can behave as selfish mobile genetic elements [75]. Thus, DIs in “*Ca.* A. scotoplanesicola” may represent mobilome-associated, selfish-element-like loci that undergo rapid gain, loss, and rearrangement. The proximity of DIs to ARs suggests a possible bacterial adaptation strategy in which the intrinsic instability of these DIs has been co-opted to generate variation in adjacent ASAP genes. In this scenario, DIs could contribute not only to protection against phages or other MGEs but also to diversification of cell-surface-associated proteins. Although no integrase or recombinase genes were annotated in the genome, several repeat-associated features were observed, including inverted repeats within DI1 and AR3, additional inverted repeats flanking an isolated ASAP gene and the 16S rRNA gene (Fig. S4), and short homologous multi-repeat sequences in intergenic regions of DI3 (Fig. 3E). These features suggest that repeat-mediated recombination may contribute to localized genome rearrangement in “*Ca.* A. scotoplanesicola”.

The body wall of *Scotoplanes* spp. consists of a thin cuticle-covered epidermis, a dermis composed of mutable collagenous tissue, and the coelomic epithelium [76], and contains microscopic calcium carbonate ossicles. The expanded ASAP repertoire may represent an adaptation to this body-wall environment and contribute to bacterial localization, persistence, or dominance. In addition, extracellular protease domains in subgroup F1 may cleave ASAPs or other surface-exposed proteins in the extracellular space, potentially altering cell-surface properties and interactions. Together, the ASAP repertoire may facilitate persistence in the subcuticular body-wall habitat and contribute to the striking dominance of this symbiont in the *Scotoplanes* body wall.

The consistent dominance of “*Ca.* A. scotoplanesicola” in the body wall across multiple *Scotoplanes* specimens raises questions about how this symbiont is maintained and transmitted among host individuals. Vertical transmission of dominant SCB has previously been proposed in the brooding brittle star *Amphipholis squamata*, where ultrastructural observations suggested bacterial transfer from the parent to brooded embryos [64,77]. Given the detection of “*Ca.* A. scotoplanesicola” signals in gonadal tissue, vertical transmission of this symbiont is a plausible hypothesis, although direct transfer from parent to embryo remains to be demonstrated. Future studies should investigate the transmission route and diversification process of “*Ca.* A. scotoplanesicola”.

Our findings reveal a previously underexplored form of echinoderm–*Mycoplasmatota* symbiosis, in which a highly reduced yet genomically plastic bacterium dominates the body-wall niche of holothurians. This study expands our view of deep-sea invertebrates as reservoirs of *Mycoplasmatota* symbioses and sheds new light on how community dominance, tissue localization, genome reduction, localized genome rearrangement, and surface-protein diversification together shape host–microbe associations.

## Acknowledgement

We thank the crew and researchers of the deep-sea cruises KS-16-18, KS-20-15 (R/V Shinsei Maru), and KH-23-5 (R/V Hakuho Maru) led by S. Kojima of the Atmosphere and Ocean Research Institute, The University of Tokyo. We also thank R. Ryusui of the Itoh Laboratory, Institute of Science Tokyo, for technical assistance with genomic library preparation. We are grateful to T. Hoshino of the Japan Agency for Marine-Earth Science and Technology (JAMSTEC) for advice on FISH probe design and sample fixation for FISH analysis, as well as for valuable comments on the manuscript.

## Declaration of generative AI and AI-assisted technologies

During the preparation of this work, the authors used ChatGPT for English language editing, and OpenAI Codex and Claude Code by Anthropic for the development, debugging, and review of analysis scripts. These tools were not used to generate scientific interpretations or conclusions. The authors independently verified the analysis scripts and results, reviewed and edited all AI-assisted content, and take full responsibility for the content of the manuscript.

## Data availability statement

The raw metagenomic sequencing reads used for genome assembly, including paired-end reads from H4, H5, and H8 and mate-pair reads from H4, will be deposited in the DDBJ Sequence Read Archive under BioProject accession number [PRJDB42313]. The 16S rRNA gene amplicon sequencing reads from St1, St2, Sk, Sh1, and Sh2 will also be deposited in the DDBJ Sequence Read Archive under the same BioProject accession number. The complete circular genome sequence of “Candidatus Abyssoplasma scotoplanesicola” BDBH4 will be deposited in DDBJ/ENA/GenBank under accession number [AP050801]. The near-full-length 16S rRNA gene sequences obtained by Sanger sequencing from H5, H8, St1, St2, Sk, Sh1, and Sh2 will be deposited in DDBJ/ENA/GenBank under accession numbers [LC934613–LC934619]. All other data supporting the findings of this study are included in the article and its supplementary material.

## Author contributions

Investigation: H.I., M.H.-T., T.T., and A.O. collected the samples; Y.Y. and H.I. performed Sanger sequencing; N.W. and W.I. performed FISH analysis. Formal analysis: Y.N., Y.G., T.I., and T.H. performed metagenomic assembly; Y.Y., Y.N., and K.T. performed genomic and gene-content analyses; Y.Y., M.H.-T., and T.T. performed amplicon sequence analysis. Funding acquisition: Y.Y. and S.Y. Visualization: Y.Y. and Y.N. Supervision: Y.N. and S.Y. Writing – original draft: Y.Y., H.I., K.T., and Y.N. Writing – review & editing: all authors. All authors approved the final manuscript and agreed to be accountable for the accuracy and integrity of the work.

---

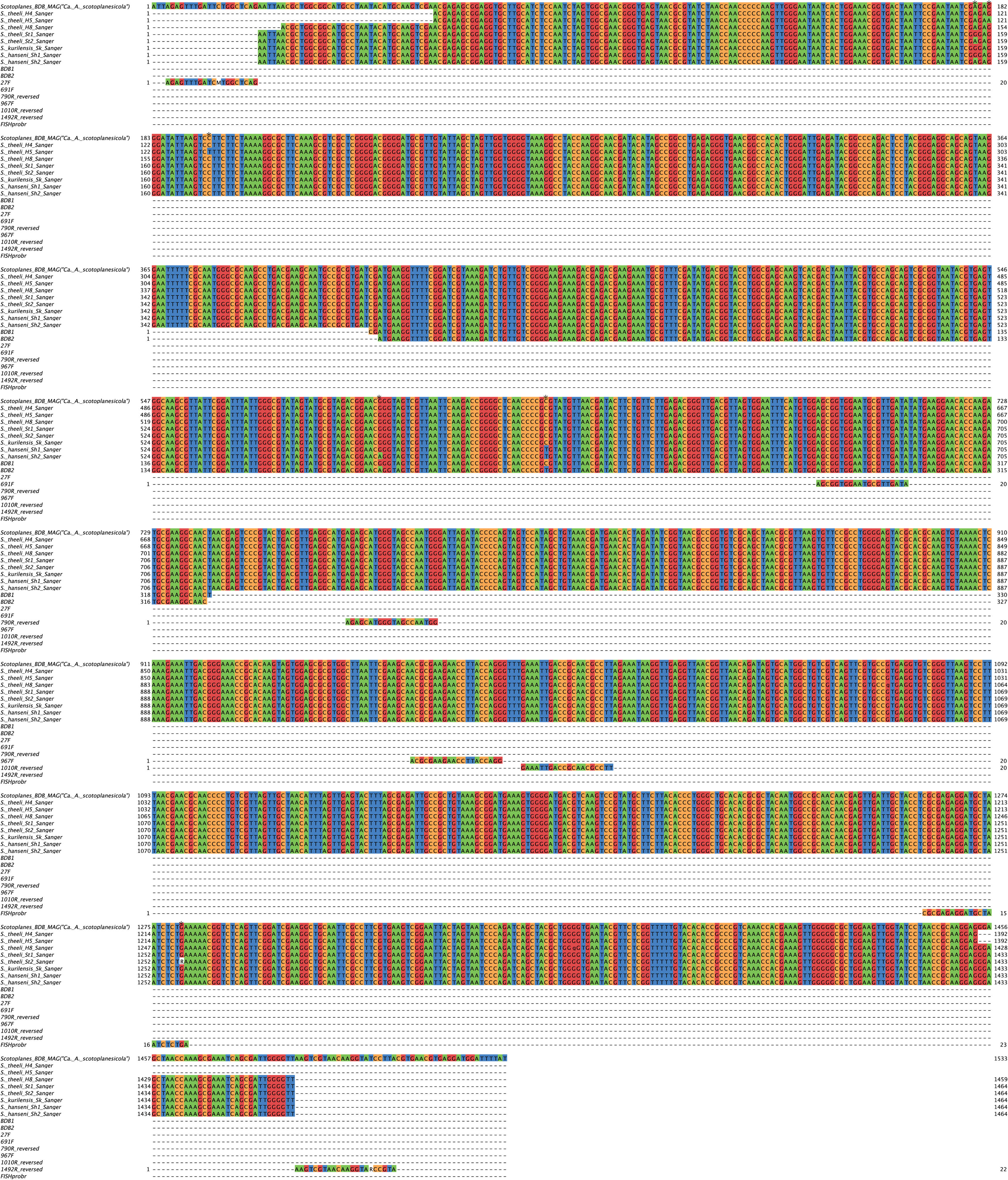

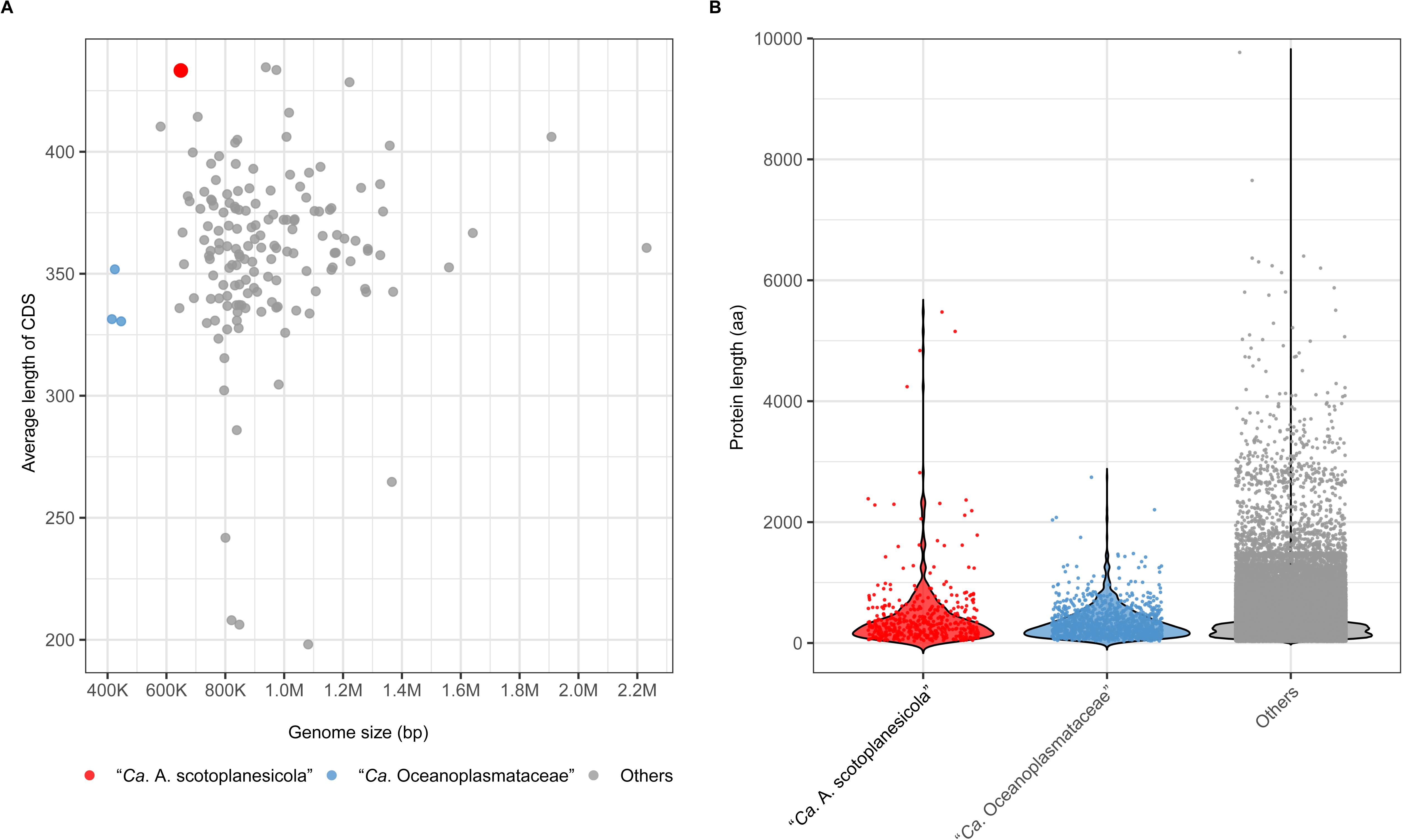

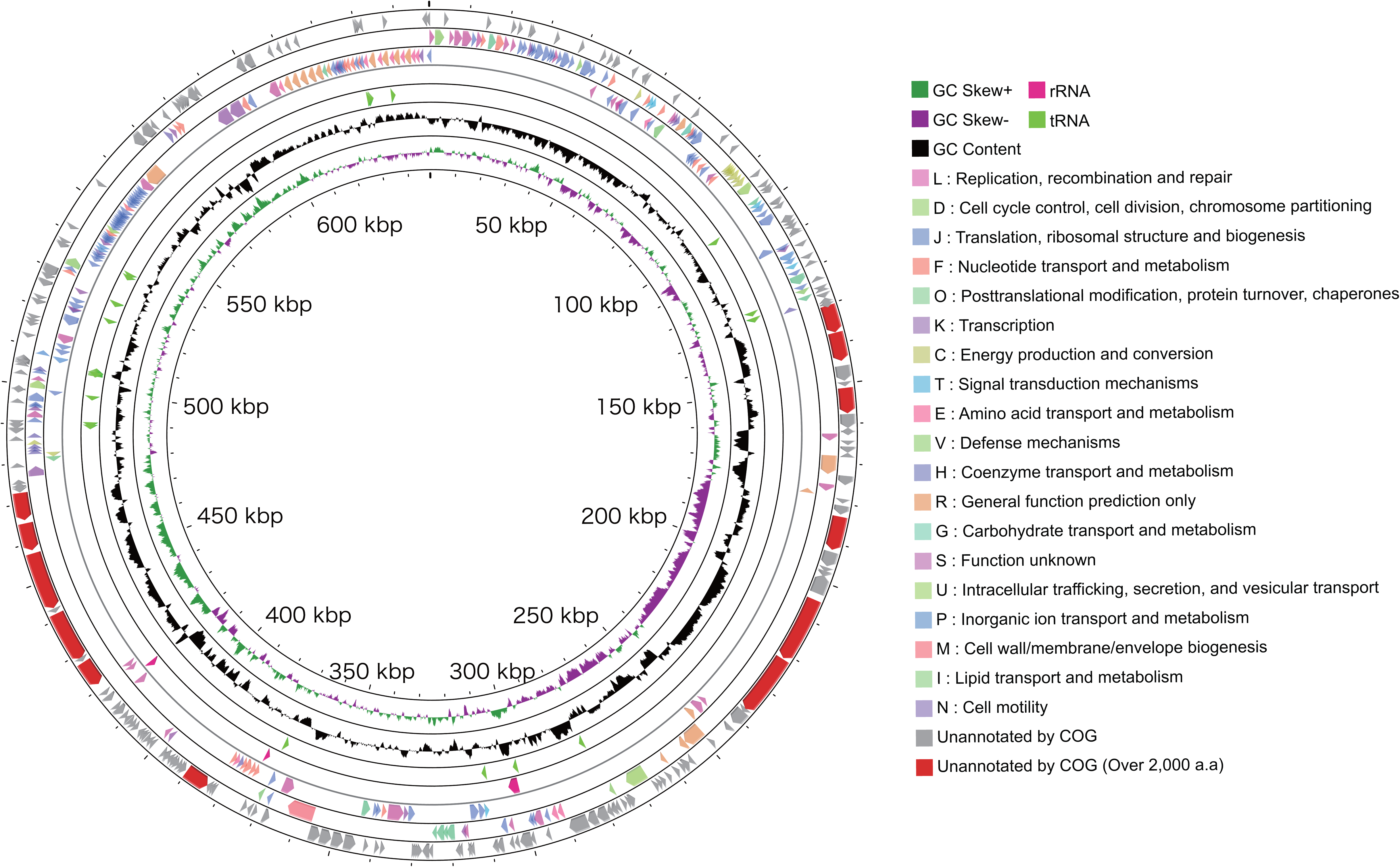

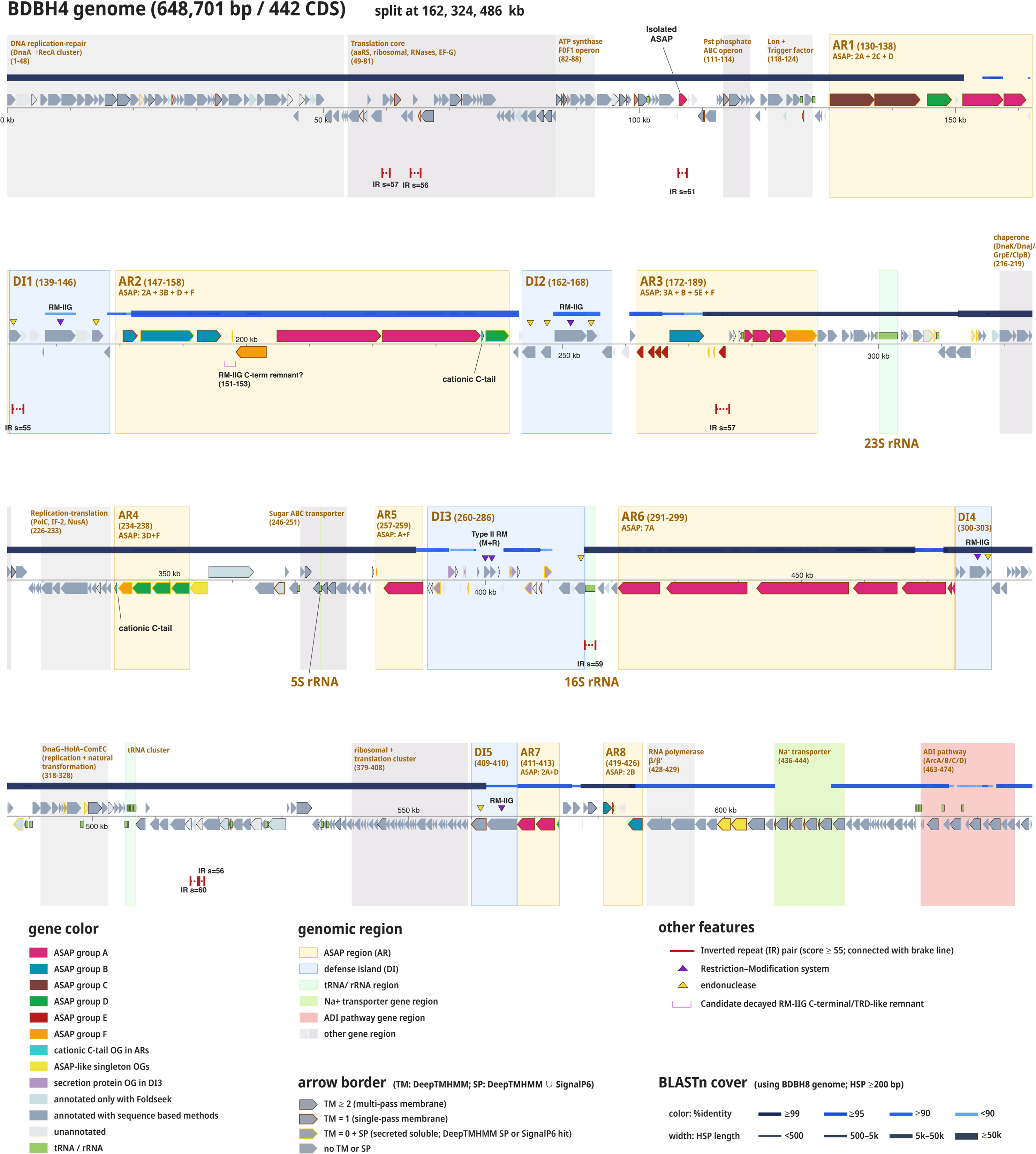

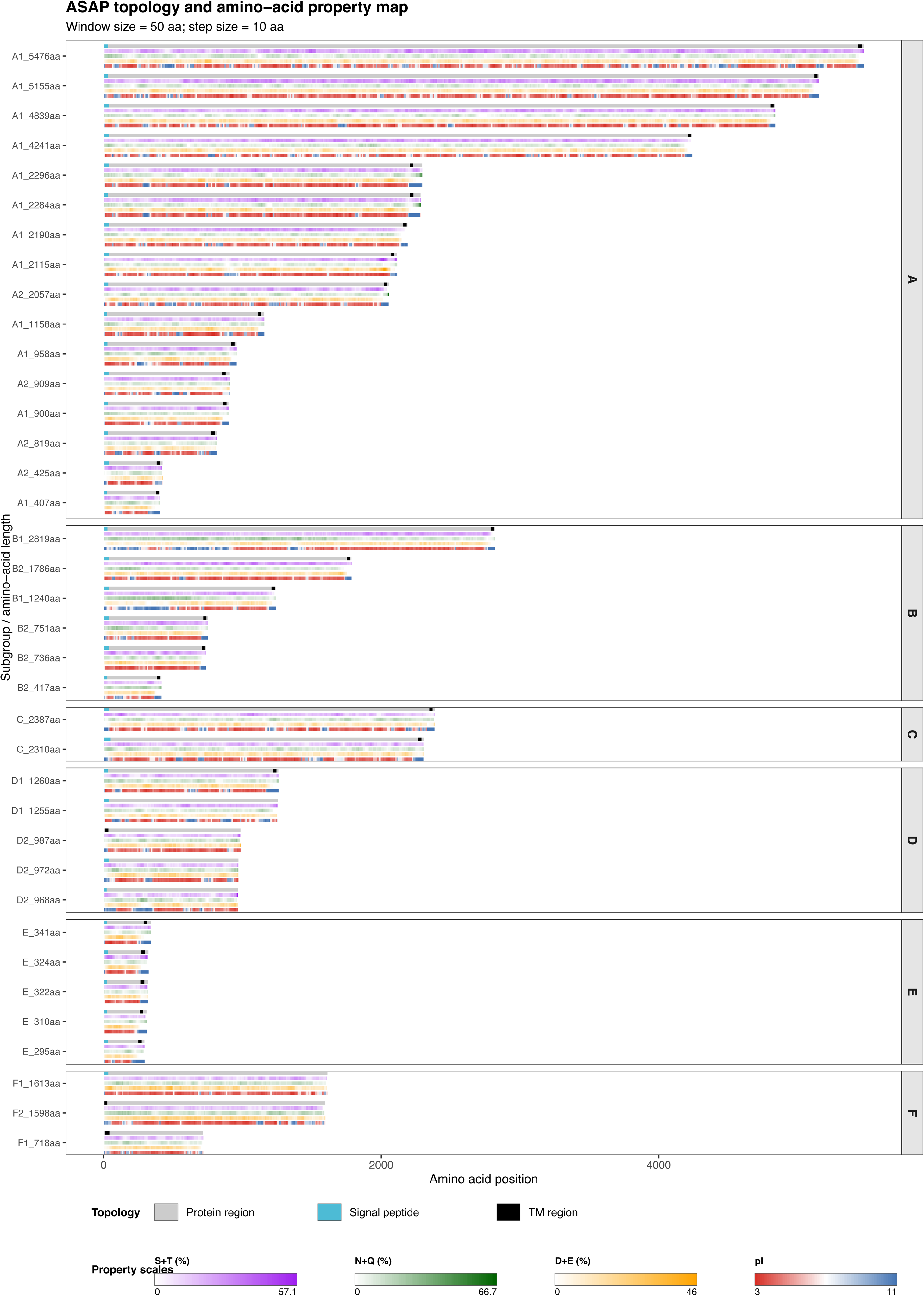

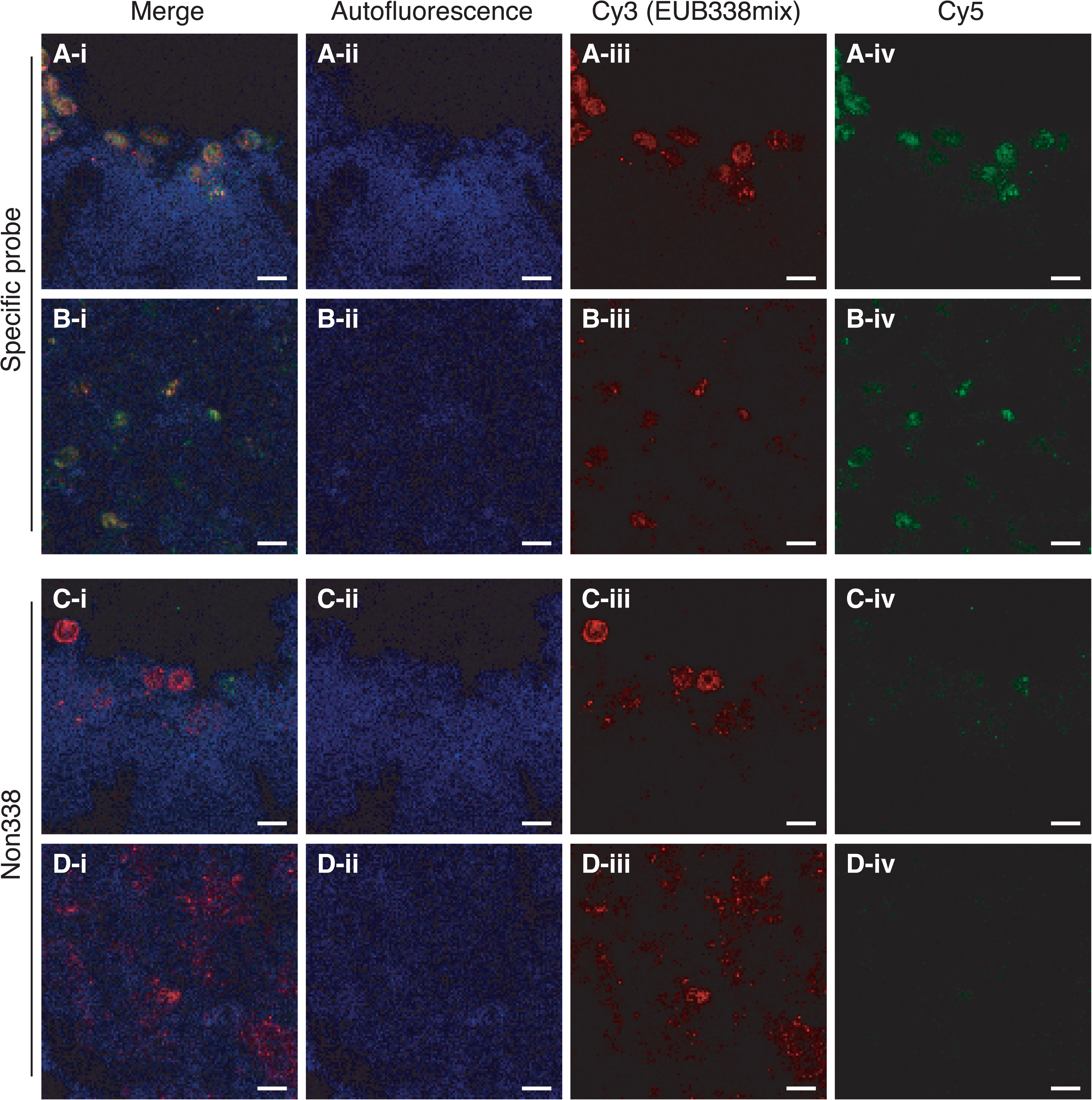

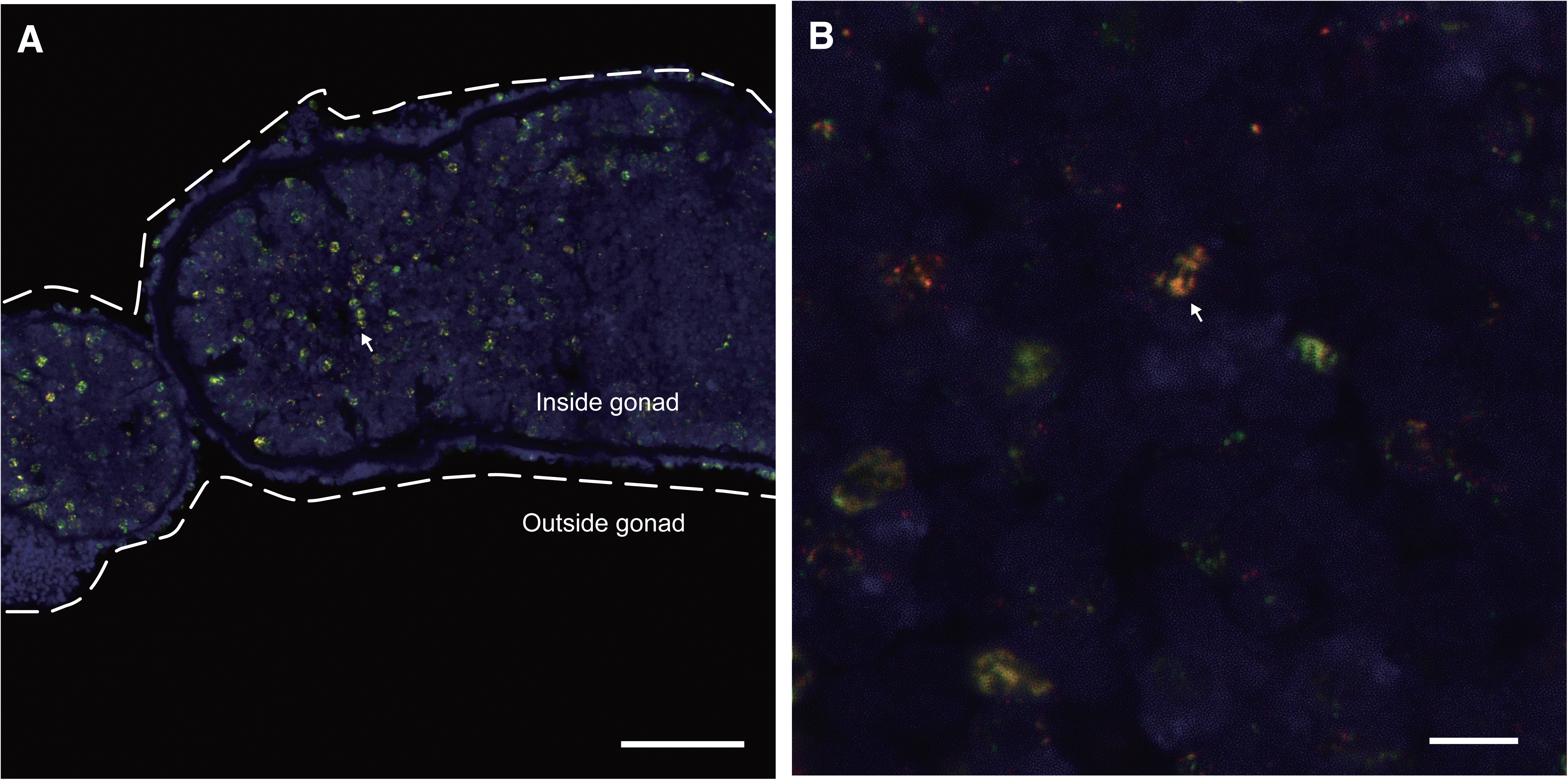

