## Supplementary Methods and Figure Legends for "Genomic, spatial, and evolutionary insights into a dominant *Mycoplasmatota* symbiont colonizing the body wall of deep-sea holothurians"

**Sanger sequencing of 16S rRNA genes**

The five *Scotoplanes* specimens (two of *S. theeli,* two of *S. hanseni,* and one of *S. kurilensis*), collected during the R/V Shinsei Maru cruise KS-20-15 and R/V Hakuho Maru cruise KH-23-5, were fixed on board in 99% ethanol overnight in a refrigerator. The ethanol was then replaced with fresh 99% ethanol, and the samples were stored at 4°C until DNA extraction. Total DNA was extracted using a DNeasy Blood and Tissue Kit (QIAGEN, Germany) according to the manufacturer's instructions.

Microbial 16S rRNA gene fragments were amplified using the universal bacterial primer set 27F/1492R [1] (Table S1). PCR amplifications were carried out in a 13 µl reaction mixture containing 1 µl of template DNA, 3.7 µl of dH2O, 0.4 µl of each primer (10 µM), and 6.5 µl of KOD One PCR Master Mix (TOYOBO, Japan). The thermocycling program consisted of an initial denaturation at 98°C for 2 min, followed by 30 cycles of 98°C for 10 s, 52°C for 20 s, and 68°C for 5 s, and a final extension at 72°C for 4 min. PCR products were purified using ExoSAP-IT (Thermo Fisher Scientific, USA). Purified PCR products were sequenced using a BigDye Terminator v3.1 Cycle Sequencing Kit (Thermo Fisher Scientific, USA). For specimens H4, H5, and H8, sequencing was performed using the internal primers 533F and 907R [2]. For the five newly collected specimens, "*Ca*. A. scotoplanesicola"-targeted primers were designed using Primer-BLAST [3] and used for sequencing, because sequencing with 533F/907R yielded reads containing many ambiguous bases. These primers were designed_primer1_691F, designed_primer2_1010R, designed_primer3_967F, and designed_primer4_790R (Table S1). Sequencing products were purified using a BigDye XTerminator Purification Kit (Thermo Fisher Scientific, USA) and analyzed using an Applied Biosystems 3500xL Genetic Analyzer (Thermo Fisher Scientific, USA).

**Structural analysis**

Structures of acidic cell-surface-associated proteins (ASAPs) were predicted using AlphaFold3 v3.0.2 [4] with customized alignments. Three alignment strategies were tested. In the first strategy, alignments were generated using the default AlphaFold3 v3.0.2 alignment pipeline. In the second strategy, group-specific alignments were generated for each ASAP group using custom homolog databases. Each database consisted of full-length BDBH4 sequences belonging to the corresponding ASAP group. For groups D and F, orthologous sequences from other "*Ca*. Oceanoplasmataceae" genomes were also included, with one and two additional sequences added to the respective databases. Alignments were constructed using MAFFT v7.526 [5] with the --auto option. In the third strategy, alignments were generated using the same homolog databases as in the second strategy, but the sequences were additionally fragmented using a 200-aa sliding window with a 100-aa step size. Terminal fragments shorter than 50 aa were discarded. This fragment-augmented database was used to improve alignment coverage for long, internally repeated proteins. Each full-length target protein was used as an independent query against the corresponding fragment-augmented database using jackhmmer [6] with three iterations. For the second and third strategies, the resulting Stockholm alignments were converted to A2M format using esl-reformat [6].

Structures were predicted using the three alignment strategies and evaluated based on mean Cα pLDDT values for 37 ASAP sequences longer than 200 aa (Table S10). For all sequences, the highest mean pLDDT was obtained using either strategy 2 or strategy 3, indicating that the default AlphaFold3 alignment pipeline was not effective for ASAPs. Strategy 3 was selected for subsequent analyses because it yielded the highest mean pLDDT for 24 of the 37 sequences. Group C showed consistently low mean pLDDT values, ranging from 37.7 to 37.8, and was excluded from subsequent structural analyses. For the remaining groups, mean pLDDT values ranged from 53.1 to 85.5 (Table S10). We considered these structures sufficiently reliable for interpreting domain-scale structural features and modular architectures.

Based on the predicted structures, 3D modular segments were defined from AlphaFold3 predicted aligned error (PAE) matrices. For each structure, the PAE matrix was first binned into 10-residue intervals. Low-confidence terminal regions were trimmed when the corresponding binned mean pLDDT was <50. The remaining region was segmented into contiguous blocks by dynamic programming, with allowed block sizes of 70–320 residues. The optimization minimized within-block mean PAE with a fixed block penalty, thereby favoring compact local modules while avoiding excessive fragmentation. For each candidate block, mean PAE was calculated from the square submatrix corresponding to all residue pairs within the block. Blocks were retained as modular segments only when their mean within-block PAE was <15 Å and their length satisfied the minimum block-size criterion.

Structural similarity among PAE-defined modular segments was quantified using Foldseek [7]. For each retained segment, a segment-specific CIF file was generated by extracting the corresponding residue range from the full-length AlphaFold3 model. All segment CIF files were converted into a Foldseek database using foldseek createdb, and all-vs-all structural searches were performed using foldseek easy-search with permissive reporting thresholds: --tmscore-threshold 0.0 -e 100 --alignment-type 1. The query TM-score (qTM) was used as the segment-pair similarity score. For each segment pair, the larger qTM value from the two search directions, max(qTM(query,target), qTM(target,query)), was used. Self-comparisons were set to 1.0, and missing Foldseek pairs were treated as 0. The resulting symmetric TM-score matrix was used to visualize structural similarity. RMSD values between PAE-defined modular segments were obtained from the same all-vs-all Foldseek alignments. For each segment pair, the RMSD from the direction with the larger qTM value was used as the representative RMSD. When both directions were available, the mean of the two directional RMSD values was also recorded as an auxiliary metric. RMSD values for self-comparisons were set to 0.

PAE-defined modular segments were clustered based on the Foldseek TM-score similarity matrix. Pairwise segment distances were calculated as 1 − TM-score similarity. Hierarchical clustering was performed in Python using SciPy (scipy.cluster.hierarchy.linkage) with average linkage on the distance matrix. Cluster membership was assigned using scipy.cluster.hierarchy.fcluster at distance cutoffs corresponding to TM-score thresholds of ≥0.75, ≥0.65, and ≥0.55. The ≥0.65 cutoff was selected for cluster identification.

Electrostatic potentials were calculated for PAE-defined modular segments and full-length AlphaFold3 models using PDB2PQR v3.6.1 and APBS v3.4.1 [8]. For segment-level calculations, the corresponding residue ranges were extracted from the full-length models and written as PDB files. Structures were converted to PQR format using PDB2PQR with the PARSE force field, retaining chain identifiers. The --neutraln and --neutralc options were used where possible to neutralize artificial termini introduced by segment extraction. No explicit pH option was supplied, and default PDB2PQR protonation settings were used. APBS input files generated by PDB2PQR were used to calculate electrostatic potentials by solving the linearized Poisson-Boltzmann equation (lpbe) with automatic multigrid focusing (mg-auto). For full-length visualizations, models were reoriented by principal component analysis of Cα coordinates before APBS calculation, and finite-difference grid dimensions were capped to keep calculations tractable. Potential maps were visualized on solvent-accessible molecular surfaces in PyMOL using a fixed red–white–blue scale of −5, 0, and +5 kT/e.

**16S rRNA amplicon sequencing**

Paired-end sequencing was performed on an Illumina MiSeq platform by Seibutsu Giken Inc (Japan). The *Scotoplanes* specimens were dissected to facilitate sampling from different body parts (1–7 body parts; Fig. 5B). These body parts were lyophilized with a VD-250 Freeze Dryer (TAITEC, Japan) and vortexed for 2 min at a speed of 1,500 rpm with Multi-Beads Shocker (Yasui Kikai Co., Japan). Each crushed sample was mixed with Lysis Solution F (Nippon Gene, Japan) and protease K (TaKaRa, Japan) and incubated for 10 min at 56 °C. After centrifugation at 12,000×g for 2 min, the supernatant was collected. DNA was purified from the supernatant using a Lab-Aid 824s DNA Extraction kit (Zeesan Biotech, China). The DNA from *S. theeli* specimens was extracted using the MPure 12 system and MPure Bacterial DNA Extraction Kit (MP Biomedicals, USA), and the DNA from *S. hanseni* and *S. kurilensis* was extracted using the Lab-Aid 824s DNA Extraction Kit (Zeesan Biotech, China). NGS libraries were prepared by 2-step tailed PCR. For the 1$^{st}$ PCR, the universal primer set amplified the v3–v4 region of the 16S rRNA gene (341F, 805R) (Table S1) [9]. For the 2$^{nd}$ PCR, sample-specific index sequences and Illumina adapter sequences were added using the primers 2ndF and 2ndR (Table S1). Sequencing was performed by Seibutsu Giken Inc. (Japan) using the MiSeq system and MiSeq Reagent Kit v3 (Illumina, USA) with 2 × 301 bp. The obtained PE sequences were processed using DADA2 and QIIME 2 for filtering (--p-trim-left-f 70 --p-trim-left-r 70 --p-trunc-len-f 250 --p-trunc-len-r 240), merging, ASV clustering, and the taxonomic assignment of the sequence [10]. The visualization was generated using QIIME 2 and Illustrator 2025.

**Fluorescence in situ hybridization (FISH)**

Body-wall tissue pieces (1 cm × 1 cm) and portions of the gonads were excised from three specimens, Sk, Sh1, and Sh2 (see Table 2), and fixed on board in 30 volumes of 4% paraformaldehyde phosphate buffer solution (Wako, Japan) at 4°C for 8–12 hours. After fixation, decant the PFA from the sample tubes, add an appropriate amount of 70% ethanol to each tube, and rinse each tube once. Add 30 times the volume of 70% ethanol to the tissue in the tube, and the tissue was stored at 4 °C until use in subsequent experiments. The remaining part of the body was preserved in the same manner as above for DNA analysis.

A new 16S rRNA-targeted probe was designed for "*Ca.* A. scotoplanesicola" using the ARB software package [11] and the SSURef_NR99_138 database from ARB SILVA (https://www.arb-silva.de/). *In silico* evaluation using the Silva SSU r138 database and the TestProbe 3.0 tool (https://www.arb-silva.de/testprobe) confirmed the probe's high specificity. Furthermore, probe optimization was validated by simulating formamide dissociation curves using mathFISH tool [12]. Initial probes included a 16S rRNA gene probe (EUB338), a designed probe, and a nonsense probe used as a negative control (NonEUB338 [13,14]) (Table S1). For hybridization, EUB338 was labeled with Cy3, whereas the SCB-designed and NonEUB338 probes were labeled with Cy5. Probes were applied in two combinations: EUB338 (Cy3) + SCB-designed (Cy5) and EUB338 (Cy3) + NonEUB338 (Cy5, negative control).

The samples stored in 70% ethanol were rinsed with 10 mM phosphate-buffered saline (PBS) three times, decalcified in a Morse solution (Wako, Japan), with the solution maintained at 4°C and exchanged approximately every 2 days until the specimen was completely decalcified. Following decalcification, the samples were rinsed in PBS three times and then sequentially dehydrated through an ethanol gradient series (70% for 1 h, 90% for 1 h, and abs. 100% for 1 h three times). The dehydrated samples were subsequently processed through a 1:1 solution of abs. 100% ethanol and toluene, two changes in toluene (1 h each), low-melting point paraffin wax (6 h, mp. 42–44°C) (Wako, Japan) and high-melting point paraffin wax (2 h, mp. 66–68°C, Wako, Japan) and embedded in the high-melting point paraffin. The paraffin-embedded samples were serially sectioned at 10 µm and collected on a MAS-coating slide (MATSUNAMI, Japan). Two serial tissue sections were dewaxed in xylene, dehydrated through 100% ethanol washes, and air-dried. The dried sections were immersed in a 20 mM Tris-HCl solution (pH 8.0) for 10 min at room temperature. To digest gram-positive bacterial cellular membranes and allow easier probe penetration into tissues, sections were incubated with a lysozyme solution (2 mg/ml in 20 mM Tris HCl solution, pH 8.0) for 15 min at 37°C and rinsed in 20 mM Tris HCl (pH 8.0) before the hybridization.

Tissue sections were covered with hybridization buffer (30% [v/v] formamide, 0.9 M NaCl, 20 mM Tris–HCl [pH 8.0], 0.01% SDS, 10% dextran sulfate, and 2.5× Denhardt's solution [Wako, Japan]) and incubated at 46°C for 30 min in a humid chamber as pre-hybridization step, and unlabeled oligonucleotide probes were added to a final concentration of 0.5 μM in the hybridization buffer. Then, the samples were incubated at 46°C for 2.5 h in the humid chamber. The sections were washed in pre-warmed washing buffer (20 mM Tris-HCl [pH 8.0], 0.112 M NaCl, 5 mM EDTA [pH 8.0], 0.01% SDS) for 30 min at 48°C to remove unhybridized probes. Amplifier probes were prepared by diluting each fluorescently labeled hairpin to 5 μM in amplification buffer (50 mM $Na_2HPO_4$, 0.9 M NaCl, 0.01% SDS, 10% dextran sulfate, 2.5x Denhardt's solution [Wako, Japan]), heating at 95°C for 90 s and cooling to 25°C for 30 min, and then mixing the hairpins to a final concentration of 1 μM each. The mixture containing the amplifier probes was applied to the sections after the washing step and incubated for 40 min at 35°C in the humid chamber. The sections were subsequently washed in the pre-cooled PBS for 10 min at 4°C to prevent probe dissociation, rinsed in ultrapure water, dehydrated in 100% ethanol, air-dried, and mounted with SlowFade Gold Antifade Mountant (Thermo Fisher Scientific, USA). Images were acquired using an Olympus FV3000 confocal microscope in super-resolution mode (count 7). Host tissue autofluorescence was excited with a 488-nm laser, and emission was collected over 500–540 nm to visualize tissue structure. Cy3 and Cy5 were excited with 561- and 640-nm lasers, respectively, and their fluorescence emissions were collected over 570–620 nm and 650–750 nm, respectively.

### Supplementary Figure legend

**Supplementary Fig. S1. Alignment of *Scotoplanes* BDB 16S rRNA gene sequences highlighting primer- and probe-binding sites (visualized in Jalview).**
Asterisks (*) mark nucleotide positions that do not perfectly match the corresponding primer/probe sequences (i.e., mismatches).

**Supplementary Figure S2. Genome size and coding sequence/protein length features of "*Ca*. A. scotoplanesicola" and related *Mycoplasmatota* genomes.**
(A) Scatter plot of genome size (bp) versus average CDS length for "*Ca*. A. scotoplanesicola", "*Ca*. Oceanoplasmataceae", and other *Mycoplasmatota* genomes. Points represent individual genomes. (B) Violin plots showing the distribution of protein lengths in each category; jittered points represent individual proteins.

**Supplementary Fig. S3. Circular genome map of the BDBH4 genome.**
The map shows CDSs on both strands, colored by COG functional category, together with rRNA and tRNA genes, GC content, and GC skew. CDSs unannotated by COG are shown separately, with unusually long hypothetical proteins (>2,000 aa) highlighted.

**Supplementary Figure S4. BDBH4 genome map.**
The circular genome of BDBH4 (648,701 bp; 442 CDSs) is shown as a linearized map split at 162, 324, and 486 kb. Predicted coding sequences are shown as arrows, with colors indicating ASAP groups, ASAP-like singleton ortholog groups (OGs), cationic C-tail OG, DI3 secretion protein OG, annotation categories, and tRNA/rRNA genes. Arrow borders indicate predicted protein topology based on DeepTMHMM and SignalP 6.0: multi-pass transmembrane proteins, single-pass transmembrane proteins, predicted secreted soluble proteins, or proteins without predicted transmembrane regions or signal peptides. Shaded regions indicate ASAP-rich regions (ARs), defense islands (DIs), tRNA/rRNA regions, the $Na^+$ transporter gene region, the ADI pathway gene region, and other conserved gene regions. Blue bars above the genome indicate BLASTn coverage against the BDBH8 genome, with bar color representing nucleotide identity and bar width representing HSP length. Inverted repeat (IR) pairs with scores ≥55 are shown with connected break lines. Positions of restriction–modification (RM) systems, endonucleases, and a candidate decayed RM-IIG C-terminal/TRD-like remnant are

also indicated.

**Supplementary Figure S5. Protein sequence maps for ASAP group members.**

Predicted transmembrane regions, signal peptide positions, serine/threonine (S/T) density, N/Q density, D/E density, and isoelectric point (pI) are shown for members of the six ASAP groups. Heatmap strips show properties calculated using 50-aa sliding windows with 10-aa steps along each protein sequence. Gray bars represent full protein lengths, light-blue segments indicate predicted signal peptide-like regions, and black segments indicate predicted transmembrane regions.

**Supplementary Figure S6. Fluorescence in situ hybridization (FISH) images of the body wall and gonads of *Scotoplanes hanseni*.**

Panels (A) and (B) show sections hybridized with the "*Ca*. A. scotoplanesicola"-designed oligonucleotide probe (green), while panels (C) and (D) show sections hybridized with the NonEUB338 negative-control probe. (A, C) Body wall sections; (B, D) gonad sections. Blue indicates host tissue autofluorescence.

**Supplementary Figure S7. Fluorescence in situ hybridization (FISH) images of the gonads of *Scotoplanes hanseni*.**

(A) and (B) show sections of the gonads. White arrows indicate aggregates of "*Ca.* A. scotoplanesicola". Images represent merged consecutive sections hybridized with the general bacterial probe EUB338 (red) and the "*Ca.* A. scotoplanesicola"-designed probe (green). Blue indicates host tissue autofluorescence. White dashed lines indicate the boundary of the gonad tissue.
